# Identification of CK2α’ selective inhibitors by the screening of an allosteric-kinase-inhibitor-like compound library

**DOI:** 10.1101/2024.01.18.576328

**Authors:** Deepti Mudaliar, Rachel H. Mansky, Angel White, Grace Baudhuin, Jon Hawkinson, Henry Wong, Michael A. Walters, Rocio Gomez-Pastor

## Abstract

Protein Kinase CK2 is a holoenzyme composed of two regulatory subunits (CK2β) and two catalytic subunits (CK2α and CK2α’). CK2 controls several cellular processes including proliferation, inflammation, and cell death. However, CK2α and CK2α’ possess different expression patterns and substrates and therefore impact each of these processes differently. Elevated CK2α participates in the development of cancer, while increased CK2α’ has been associated with neurodegeneration, especially Huntington’s disease (HD). HD is a fatal disease for which no effective therapies are available. Genetic deletion of CK2α’ in HD mouse models has ameliorated neurodegeneration. Therefore, pharmacological inhibition of CK2α’ presents a promising therapeutic strategy for treating HD. However, current CK2 inhibitors are unable to discriminate between CK2α and CK2α’ due to their high structural homology, especially in the targeted ATP binding site. Using computational analyses, we found a potential Type IV (“D” pocket) allosteric site on CK2α’ that contained different residues than CK2α and was distal from the ATP binding pocket featured in both kinases. With this potential allosteric site in mind, we screened a commercial library containing ∼29,000 allosteric-kinase-inhibitor-like compounds using a CK2α’ activity-dependent ADP-Glo^TM^ Kinase assay. Obtained hits were counter-screened against CK2α revealing two CK2α’ selective compounds. These two compounds might serve as the basis for further medicinal chemistry optimization for the potential treatment of HD.

## Introduction

Human protein kinase CK2 (formerly known as Casein Kinase 2) is historically the first kinase ever reported [1]. CK2 is a constitutively active serine/threonine kinase that belongs to the CMGC family (<u>C</u>DKs, <u>M</u>AP kinases, <u>G</u>SKs, and <u>C</u>DK-like kinases) and controls a multitude of cellular processes such as cell growth, proliferation, differentiation, inflammation, and cell death [2, 3]. It is not surprising that dysregulation of CK2 contributes to the etiology of multiple human diseases, including cancer, neurodegeneration, and even COVID-19 viral infection [4–6]. Therefore, CK2 has been the subject of extensive research for the development of effective and selective inhibitors for the treatment of these diseases, with a special focus on cancer treatment [6, 7]. However, despite decades of research since the first CK2 inhibitor was identified in 1990, only silmitasertib (CX-4945) has received FDA approval with limited use in clinical trials for some forms of cancer [8–12]. Recent studies in cellular and animal models have demonstrated that CK2 inhibition is also a promising therapeutic strategy for the treatment of neurodegenerative diseases like Alzheimer’s (AD), Parkinson’s (PD), and Huntington’s disease (HD) [13–18], but CK2 pharmacotherapy research in this field has been limited.

CK2 is a holoenzyme that contains two catalytic subunits, CK2 alpha (CK2α) and/or CK2 alpha prime (CK2α’) that form a tetrameric complex with a CK2β dimer, a non-catalytic scaffolding subunit. Both catalytic forms are active in the absence of CK2β, although CK2β can confer substrate specificity [19–21]. In contrast to CK2α, which is an essential protein ubiquitously expressed with hundreds of substrates, CK2α’ is dispensable since KO mice are viable, and its expression is limited to brain and testes with only a few substrates assigned to date [2, 3, 22, 23]. CK2α and CK2α’ share high structural homology, but their abundance, tissue distribution, and substrate targets suggest discrete and specialized biological functions [21, 24–26]. This is supported by studies using specific conditional deletion of CK2α or CK2α’ in neurons within the mouse striatum, a brain region that controls movement and some forms of cognition, that resulted in differential effects in locomotion, exploratory behavior, and motor learning [27]. Therefore, selective inhibition of the CK2 catalytic subunits might result in different biological outcomes.

Due to the enhanced abundance of CK2α and its specific upregulation in cancer [26, 28, 29], this subunit has received significantly more attention than CK2α’ in phenotypic and pharmacological studies, and minimal studies have addressed potential subunit-specific dysregulation or contribution in different diseases in which altered CK2 activity has been reported. Our recent work has highlighted the importance of conducting biological studies where the two subunits are investigated. We have demonstrated that CK2α’, but not CK2α or CK2β, is inappropriately upregulated in HD and found that CK2α’ is responsible for the phosphorylation of key substrates that contribute to aggravating HD pathology [30, 31]. Genetic studies knocking out CK2α’, but not CK2α, improved HD-related phenotypes, supporting CK2α’ specific inhibition as a potential therapeutic strategy in HD [30, 32].

Increased CK2 activity in affected tissues of other neurodegenerative diseases like AD and PD, contribute to the reduction of neuronal survival and worsening neurodegeneration [16, 33–37] although whether CK2α’ isoform is preferentially induced in these NDs is still unresolved. These studies suggest that access to selective probes for the CK2α’ subunit would provide essential tools for investigating its role in neuropathological processes.

Several approaches have been used to develop CK2 inhibitors [7]. These approaches include blocking the substrate channel, disrupting the holoenzyme assembly by interfering with CK2β binding, and targeting putative allosteric sites outside the catalytic box [38–40], although the most studied mode of inhibition has been through ATP analogs that compete with ATP to prevent substrate phosphorylation [8, 39, 41–43]. The ATP binding site of CK2α and CK2α’ is formed between the N– and C-terminal domains linked *via* an alpha helix (αD) loop, and share high homology [2, 44]. Due to the overall sequence similarity between CK2α and CK2α’ we felt that the best strategy to selectively target these subunits was to identify novel and specific allosteric inhibitors. To date, three well-elucidated major allosteric sites in CK2 have been the target for the development of allosteric inhibitors: the α/β interface (Site 1) [40], the αD pocket (Site 2) and the interface between the αC helix and the glycine-rich loop (G– loop or Site 3) [45] although no information regarding the degree of specificity of these compounds towards CK2α or CK2α’ has been reported. On the other hand, a recent structural study has revealed that most of the previously proposed allosteric CK2 inhibitors might indeed act through the ATP site [46], questioning the specificity of these inhibitors over other kinases. The CK2α subunit presents 20 additional amino acids at the C-terminus absent in CK2α’ along with discrete amino acid sequence variations within the ATP site that could confer different binding properties to ATP analogs [2].

Developing selective inhibitors presents a significant challenge, however when successful, this would open a new avenue for developing disease-modifying therapeutics for different human diseases. Only a few inhibitors have been reported as ATP-site Type I and Type II selective inhibitors at CK2 [47–49]. We felt another strategy was required to expand the discovery of CK2α’ selective inhibitors targeting an allosteric pocket outside the conserved, orthosteric ATP site. This strategy has provided drugs against other kinases that combat the drawbacks of ATP-site inhibitors, which typically feature limited selectivity and the potential for developing drug resistance. Using CASTp we found a potential Type IV (“D” pocket)[45] allosteric site on CK2α’ that contained different residues than CK2α and was distal (N-terminus) from the more frequently targeted ATP binding pocket featured in both kinases. We felt this site might provide a better chance for CK2α’ selectivity than the previously targeted sites, because of the difference in amino acids lining this putative binding pocket. With this promising potential binding pocket as our target, we performed a screen of a commercial library of ∼29,000 allosteric kinase inhibitor-like compounds (The ChemDiv Allosteric Kinase Inhibitor (CDAKI) Library) using a CK2α’ activity-dependent ADP-Glo^TM^ Kinase assay. The resulting hits were counter-screened against CK2α, revealing two selective compounds that were further validated using an orthogonal radioligand-based assay.

## Results and Discussion

### CK2α’-dependent ADP-Glo^TM^ assay development and optimization

CK2α and CK2α’ possess different substrates, and contribute differently to various human diseases. Therefore, it is necessary to develop specific CK2 inhibitors that can discriminate between these two catalytic subunits. Due to the relevance of CK2α’ in the pathophysiology of HD, the strong effects of genetic ablation of this subunit in the amelioration of HD-like phenotypes in rodents, and the lack of effective therapies for HD, we have focused our work on the identification of potential CK2α’ selective inhibitors.

Using CASTp (2) and PDB ID 3e3b, we determined that CK2α’ did possess a potential binding site outside the ATP-binding pocket (Pocket 2) that differed in the amino acid sequence of CK2α (**Fig. 1A, B**). This putative allosteric site was in the same region where other Type IV allosteric kinase inhibitors were known to bind [45]. To identify new CK2α’ hits that could potentially target this Type IV allosteric site, we decided to screen a commercial library of 28,812 allosteric-kinase-inhibitor-like compounds (The ChemDiv Allosteric Kinase Inhibitor (CDAKI) Library). This library is a focused collection of compounds from the ChemDiv compound collection, structurally based on known allosteric kinase inhibitors and their analogues. It is enriched in lead-and drug-like molecules with favorable physicochemical properties and low numbers of PAINS [50]. We chose to screen these compounds using the HTS-tested ADP-Glo^TM^ Kinase assay [51] for its ability to directly measure kinase enzyme activity as it relies on the detection of ADP generated from ATP used in the catalytic reaction (**Fig. 2A**). To determine the limit of detection of ADP produced, an ATP to ADP standard curve was generated. The assay was able to detect ATP conversion as low as 1% with a signal-to-background ratio of 2:1. A 10% ATP conversion, at which most enzyme reactions are performed, yielded an excellent signal-to-background ratio of 20:1 (**Supplementary** Fig. 1A).

**Figure 1.**
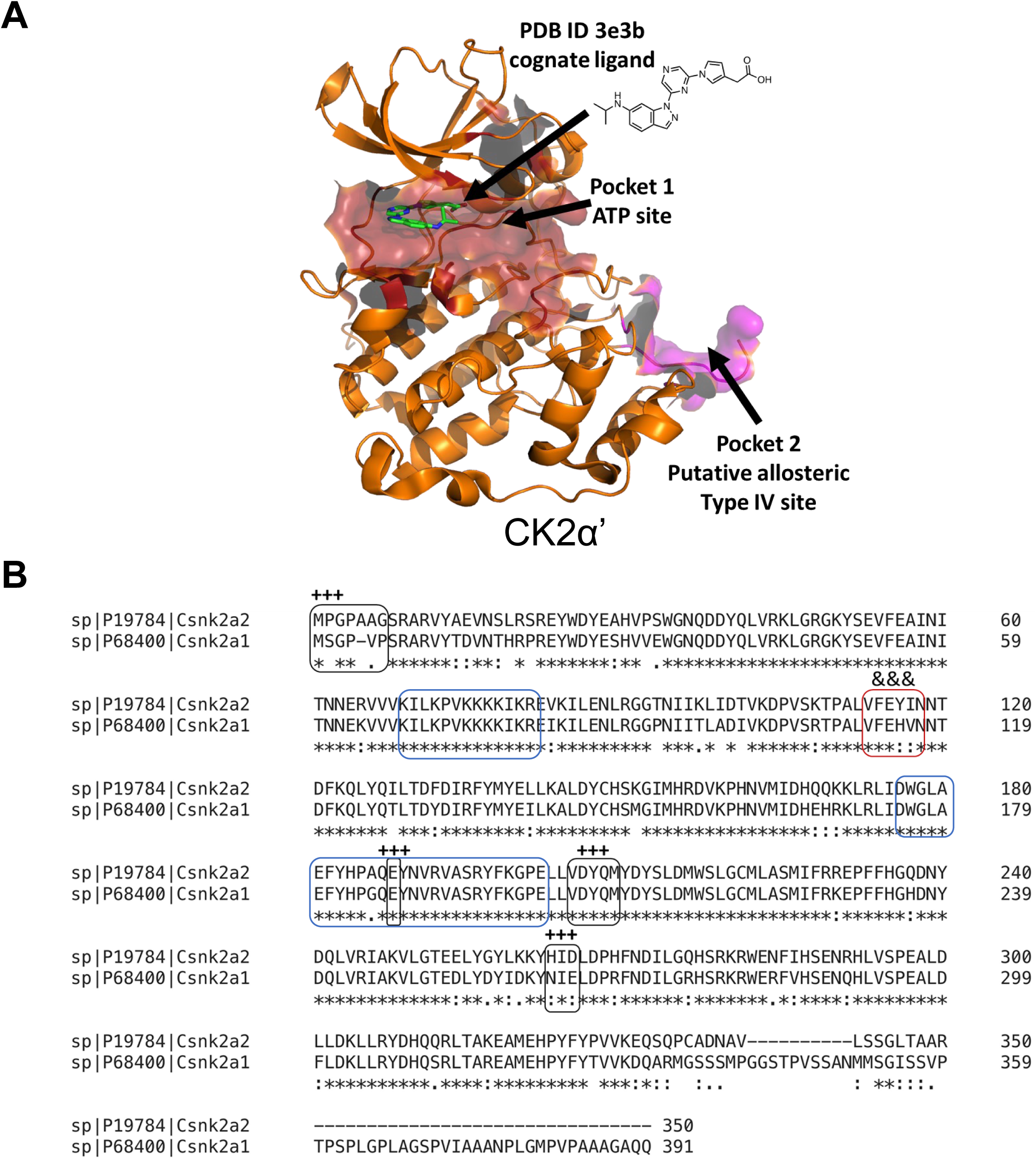
(**A**) CK2α’ protein structure indicating a putative Type IV binding pocket as computed using CASTp and visualized using PyMOL. **(B)** Protein sequence alignment for human CK2α (Csnk2a1) and CK2α’ (Csnk2a2). The black boxes (+++) denote a putative allosteric site Type IV. The red box (&&&) denotes kinase hinge region and the blue boxes denote the ATP binding site.

**Figure 2.**
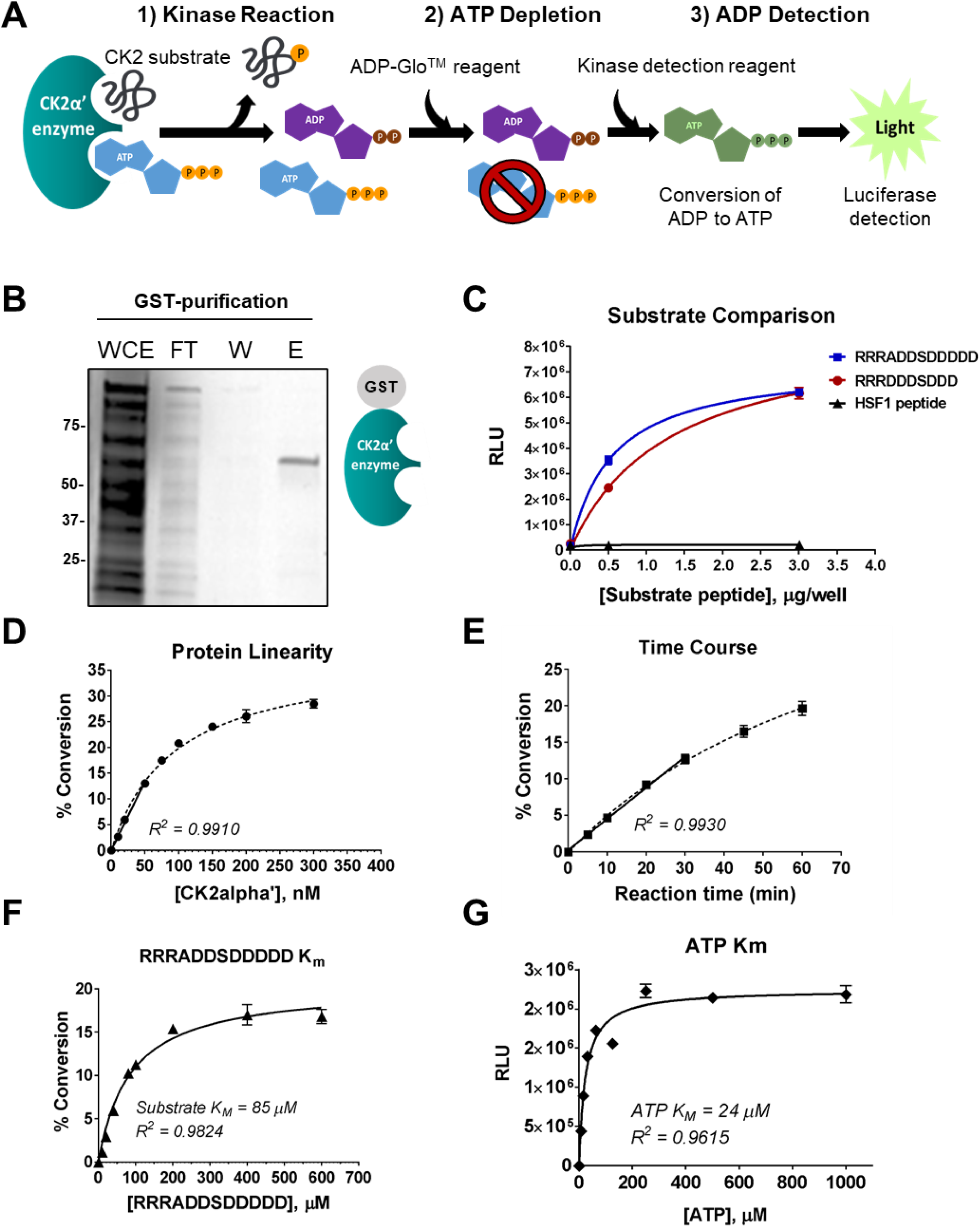
CK2α’-dependent ADP-Glo^TM^ assay development and optimization. (A) The ADP-Glo^TM^ Kinase assay is performed in three steps: 1) the target kinase phosphorylates a specific peptide substrate generating ADP from the provided ATP, 2) the kinase reaction is terminated by addition of the ADP-Glo^TM^ reagent, and the residual ATP is depleted, and 3) the ADP is converted into newly synthesized ATP by the addition of a kinase detection reagent. The newly synthesized ATP is converted to light using the luciferase/luciferin reaction. The luminescence generated is proportional to the amount of ADP generated in the kinase reaction, which is indicative of the proportional kinase activity. (B) CK2α’ enzyme (GST-tagged) was purified from bacterial cultures using a Glutathione Affinity purification system. (C) Three peptides were tested as possible substrates for CK2α’. PPQSPRVEEASP, a 12-mer polypeptide of HSF1 for which previous CK2α’-dependent phosphorylation has been reported, failed to show any activity. Of the two peptides that showed excellent activity, the commercial CK2 synthetic peptide (RRRADDSDDDDD) was selected as the final substrate for the HTS standard assay. (D) To determine initial velocity conditions, a reaction containing 80 µM of CK2 peptide substrate and 200 μM ATP with increasing concentration of recombinant human GST-CK2α’ enzyme (10 nM to 300 nM) was allowed to proceed at RT for 30 minutes. An enzyme concentration of 40 nM, at which 10% of the substrate has been converted into product, was selected for the standard assay to remain in the linear regime. (E) The activity of 40 nM CK2α’ was measured at 5, 10, 20, 30, 45 and 60 min to determine the optimal reaction time to retain linearity. The 30 min time point was selected, as it showed good linearity while maintaining a percent conversion of 10 – 15%. (F) A reaction containing 40 nM recombinant human GST-CK2α’ enzyme and 200 μM ATP with increasing concentrations of CK2 peptide substrate (0 – 800 μΜ) was allowed to proceed at RT for 30 minutes. The data was fitted to the Michaelis-Menten equation to obtain the indicated substrate Km value. (G) A reaction containing 20 nM recombinant human GST-CK2α’ enzyme and a saturating concentration (600 μM) of CK2 peptide substrate, with increasing concentrations of ATP (7.8 μΜ – 1 mM) was allowed to proceed at RT for 30 minutes. The data was fitted to the Michaelis-Menten equation to obtain the indicated ATP Km value.

The minimum consensus sequence for CK2 substrate recognition has been defined as S/T-X-X-E/D/pX, often with acidic residues also in position n+1 and/or n+2 [21, 52]. CK2α’ enzyme (GST-tagged) was purified from bacterial cultures using a glutathione affinity purification system (**Fig. 2B**), and we tested three possible peptides as CK2α’ kinase substrates. We used a custom CK2 peptide (RRRDDDSDDD) [53] and a commercial CK2 synthetic peptide (RRRADDSDDDDD) [54], both previously used as CK2 substrate peptides in previous analyses. We also used PPQSPRVEEASP, which, although it deviates from the canonical CK2 substrate binding sequence, represents a 12-mer polypeptide of a transcription factor (HSF1) previously shown to be phosphorylated by CK2α’ [30]. However, PPQSPRVEEASP did not show any activity in preliminary ADP Glo^TM^ activity assays. In contrast, RRRDDDSDDD and RRRADDSDDDDD both showed excellent and comparable levels of activity (**Fig. 2C**). The synthetic peptide RRRADDSDDDDD was selected as the final substrate for the HTS standard assay, based on its availability as a commercial product.

We next determined the initial velocity conditions of the enzymatic reaction which is necessary to further determine the inhibitory potency of test compounds towards the target enzyme. The initial velocity is the best measure of enzyme reaction rate, and the reaction is most sensitive to the effect of reversible inhibitors in this phase [55]. To experimentally determine the initial velocity, or the linear portion of the enzyme reaction where less than 10 –15% of the substrate has been depleted, we recorded an enzyme reaction progress curve. By choosing a fixed reaction time, reaction temperature, and substrate concentration, the enzyme activity in terms of percent conversion was measured by varying the enzyme concentration (**Fig. 2D**).

The assay was linear up to 13% conversion at an enzyme concentration of 50 nM. To ensure that the reaction remained in the linear regime, a concentration of 40 nM was selected for the CK2α’ enzyme. Next, we examined the activity of 40 nM CK2α’ at different time points to determine the optimal reaction time to retain linearity (**Fig. 2E**). A reaction time of 30 min showed good linearity, while maintaining a percent conversion of 10 – 15%. Hence, the 30 min time point was selected for subsequent experiments. Once these initial velocity conditions were established, a substrate saturation curve was recorded to determine K_M_ of the CK2 peptide substrate. To afford the ability to identify compounds of all inhibition modalities, the substrate concentration was chosen to equal its K_M_ value [55]. Hence, a substrate concentration of 80 µM was selected for the standard HTS assay (**Fig. 2F**). Next, the K_M_ for ATP was determined using a saturating concentration of the substrate (**Fig. 2G**). Using a high concentration of ATP (∼8X K_M_) would reduce the potency of ATP-competitive inhibitors, thereby biasing the assay towards detecting allosteric inhibitors [51]. Hence, an ATP concentration of 200 µM was chosen for our standard HTS assay.

In addition to the parameters described above, we also optimized the composition of the enzyme reaction buffer using the recommended guidelines from the ADP Glo^TM^ kit as a starting point. The effect of ionic salts (NaCl), BSA, and detergents (Brij 65, Triton X100) was also evaluated (**Supplementary** Fig. 1B-D). While adding 100 mM NaCl caused a modest increase in enzyme activity, the addition of 0.01% detergent resulted in a 5-fold increase in percent conversion. Therefore, the final optimized assay buffer contained 50 mM Tris-HCl, pH 7.5, 10 mM MgCl2, 0.1 mM EDTA, 100 mM NaCl, 2 mM DTT, 0.01% Triton X-100. Finally, the tolerance of the assay to DMSO was studied, as our compound libraries are dissolved and stored in 100% DMSO. The assay showed no detrimental effect of DMSO for up to 5%.

### High throughput screen and counter screen identified two CK2**α**’ selective hits

The optimized ADP Glo^TM^ standard HTS assay was used to screen 28,812 compounds of the ChemDiv Allosteric Kinase Inhibitor (CDAKI) Library (ChemDiv, Inc) (**Fig. 3**). The reference inhibitor silmitasertib (CX-4945) was included as a positive control on every screening plate. Vertical validation experiments using alternating columns of no inhibitor, no enzyme, and an IC_50_ concentration of CX-4945 as a positive control yielded an average Z’ of 0.85 ± 0.029, which is substantially higher than the recommended minimum value of 0.5 for HTS. A pilot screen comprising the TOCRIS kinase inhibitor toolbox (80 compounds) and the GSK kinase inhibitor PKIS2 collection (473 compounds) produced a similar pattern of apparent inhibitors on a 3X repeat test, indicating that the assay was reproducible. Of the 28,812 compounds screened, 105 that produced > 50% inhibition (0.4% hit rate) were selected as screening hits. 95 were confirmed as inhibitors (with IC_50_ < 100 µM) in dose-response experiments conducted with cherry-picked compounds from DMSO stocks. 65 prioritized hits (with IC_50_ < 10 µM) were further tested in an interference assay to identify luciferase-inhibiting compounds. None of these compounds interfered with the ADP Glo^TM^ detection reagents (**Supplementary Table 1**).

**Figure 3.**
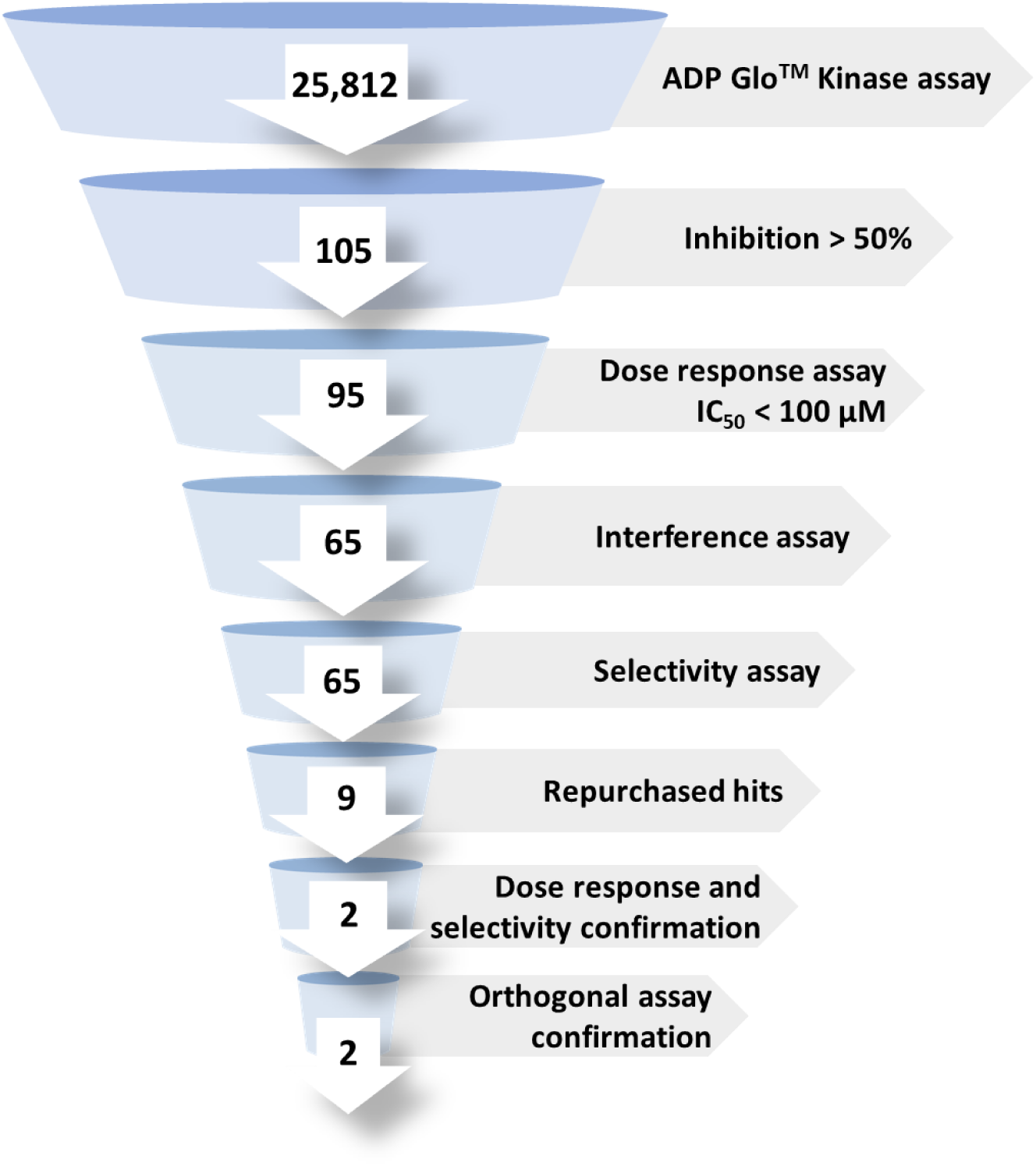
Flowchart for the discovery of novel CK2α’ selective inhibitors.

The 65 confirmed active hits were counter-screened along with CX-4945 using CK2α as the target kinase. Hits were tested in a dose-response assay to evaluate inhibitor selectivity for CK2α’ over CK2α. Nine compounds that showed >1.5-fold selectivity were repurchased for selectivity confirmation. Liquid chromatography-mass spectrometry (LC-MS) and nuclear magnetic resonance (NMR) spectroscopy confirmed compound identity and purity greater than 92% for all 9 compounds (Supplementary Fig. 2-10). CX-4945, along with seven repurchased hits (compounds **3-9**), showed no selectivity between CK2α’ and CK2α (**Fig. 4A-C**, **Table 1, Supplementary** Fig. 11). An IC_50_ of 8-10 nM for CX-4945 was obtained, which is comparable with values reported in the literature (IC_50_ of 1 nM in a cell-free assay) [12]. Two of the repurchased compounds (**1** and **2**) showed enhanced selectivity for CK2α’ over CK2α (**Fig. 4A, D, E**, **Table 1**). Compound **1** showed a 5-fold selectivity for CK2α’ over CK2α, and a 30% higher efficacy. Compound **2** showed a 2-fold selectivity and 2-fold improvement of efficacy for CK2α’ over CK2α. In addition, we confirmed that compounds **1** and **2** presented similar IC_50_ between the CK2α’-GST enzyme (produced in-house and used in the HTS) and a commercially obtained CK2α’ enzyme tagged with His (CK2α’-His) demonstrating that the GST tag used in the in-house produced CK2α’ was not introducing confounding selectivity (**Supplementary** Fig. 12).

**Figure 4.**
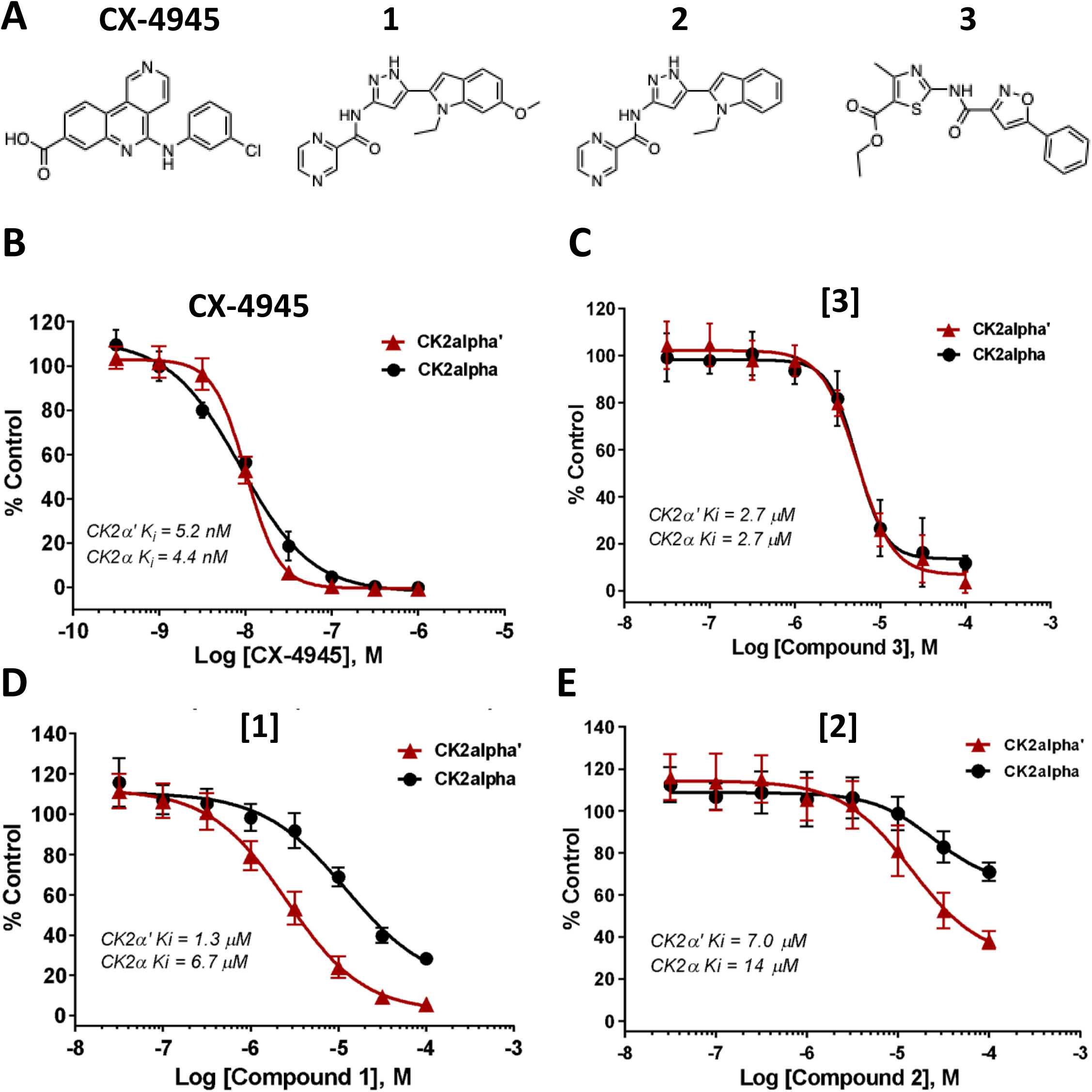
Inhibitor selectivity for CK2α’ over CK2α. (A) Structures of the reference inhibitor silmitasertib (CX-4945) and 3 hits identified in the HTS campaign. (B) CX-4945, a potent ATP-competitive inhibitor of CK2, showed no selectivity between CK2α’ and CK2α. (C) Compound 3 is 1 of 7 repurchased hits that showed no selectivity between CK2α’ and CK2α. (D) Compound 1 showed a 5-fold selectivity for CK2α’ over CK2α, and a 30% higher efficacy. (E) Compound 2 showed a 2-fold selectivity for CK2α’ over CK2α, and a 2-fold improvement of efficacy.

**Table 1:** Summary of kinase profiling of CX-4945, 1 and 2 tested against CK2α and CK2α’ in the Promega ADP-Glo Activity Assay.

| Compound ID: | Compound IC<sub>50</sub> (μM) | | Fold change:<br>$\alpha'$ vs $\alpha$ |
| --- | --- | --- | --- |
| | CK2α | CK2$\alpha'$ | |
| **CX-4945** | $0.011 \pm 0.00041$ | $0.010 \pm 0.00088$ | 1.1 |
| **1** | $15 \pm 0.037$ | $2.6 \pm 0.0022$ | 5.8 |
| **2** | $49 \pm 0.29$ | $15 \pm 0.021$ | 3.3 |

The selectivity of **1** and **2** was further confirmed in the Kinase HotSpot orthogonal assay (Reaction Biology, Malvern, PA) (**Fig. 5A**) using CK2α’, CK2α and another important kinase GSK3β, also expressed in the brain [56]. The HotSpot assay uses ^33^P-ATP in the kinase reaction and measures kinase activity using a P81 filter-binding method [57]. CX-4945 showed increased selectivity for CK2α’ and CK2α compared to GSK3β, with comparable IC_50_ between CK2α’ and CK2α (**Fig. 5B**, **Table 2**), as observed using the ADP-Glo^TM^ assay (**Fig. 4B**). For this assay, the IC_50_ for **1** and **2** were one order of magnitude smaller than those obtained using the ADP-Glo^TM^ assay, perhaps due to the increased sensitivity of The HotSpot assay (**Tables 1, 2**). Selectivity of **1** and **2** for CK2α’ was also confirmed, although **2** also showed a greater inhibition of GSK3β compared to **1**(**Fig. 5C, D**).

**Figure 5.**
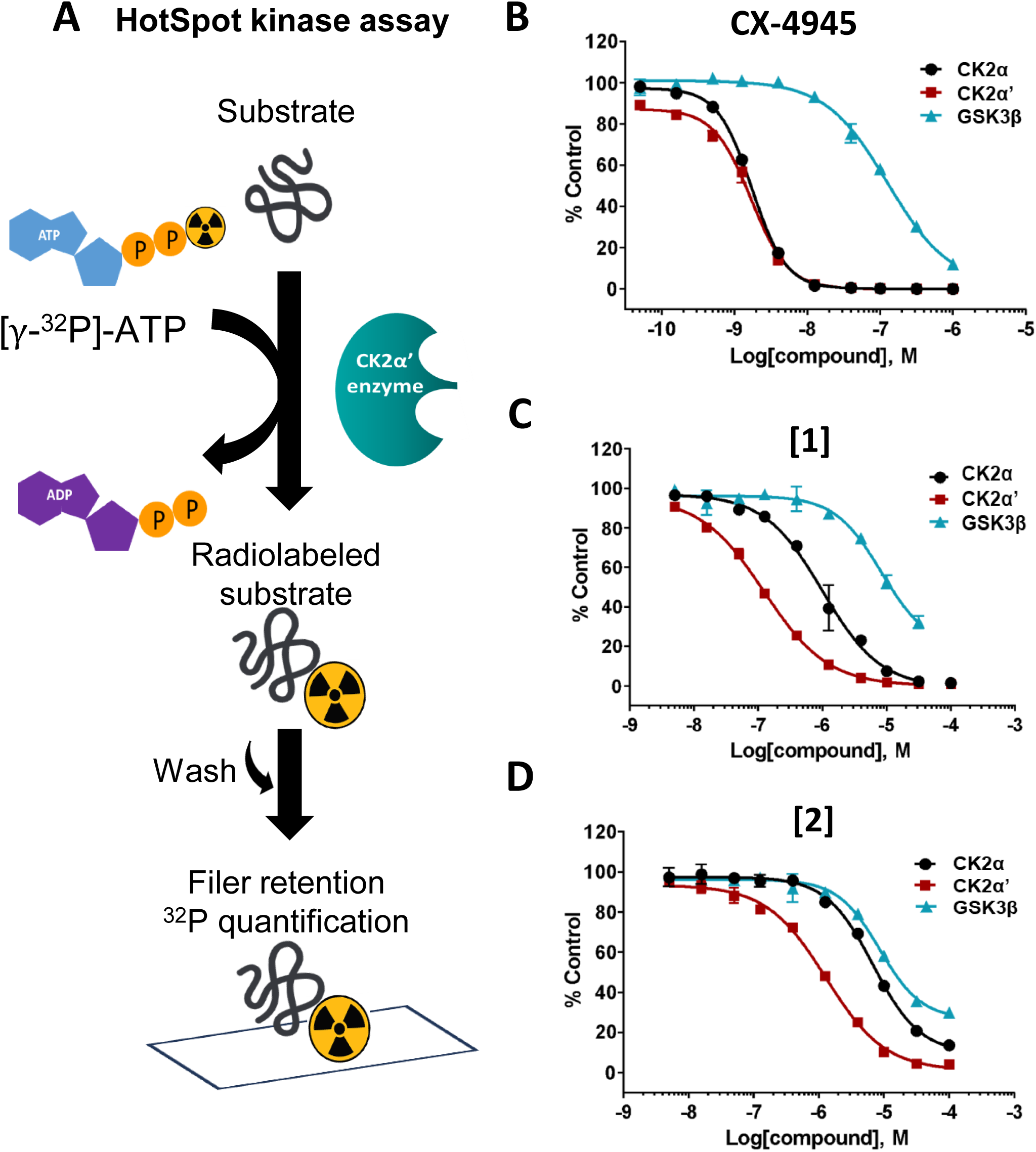
Orthogonal assay confirmation of hit selectivity. (A) The Kinase HotSpot orthogonal assay measures kinase activity based on the transfer of ^33^P-labelled phosphate from ATP to the kinase substrate. (B) CX-4945 showed increased selectivity for CK2α’ and CK2α compared to GSK3β, with comparable IC_50_ between CK2α’ and CK2α in the orthogonal assay. (C,D) Selectivity of compounds 1 and 2 for CK2α’ over CK2α was confirmed, with compound 2 showing a greater inhibition of GSK3β compared to compound 1.

**Table 2:** Summary of kinase profiling of CX-4945, 1 and 2 tested against CK2α, CK2α’ and GSK3β in the Kinase HotSpot Assay.

| Compound ID: | Compound IC<sub>50</sub> (μM): | | | Fold change:<br>$\alpha'$ vs $\alpha$ | Fold change:<br>$\alpha'$ vs GSK3β |
| --- | --- | --- | --- | --- | --- |
| | CK2α | CK2$\alpha'$ | GSK3β | | |
| **CX-4945** | 0.0019 ± 0.000011 | 0.0017 ± 0.00013 | 0.14 ± 0.013 | 1.1 | 82 |
| **1** | 0.97 ± 0.14 | 0.11 ± 0.00035 | 8.6 ± 0.68 | 8.8 | 78 |
| **2** | 9.1 ± 0.39 | 1.2 ± 0.026 | 21 ± 0.44 | 7.6 | 17.5 |

### CK2**α**’ allosteric selective hits compete with ATP

Finally, since CK2α’ and CK2α share a high structural homology, it was imperative to identify small-molecule scaffolds that are allosteric inhibitors of CK2α’ that do not bind to the ATP-binding site of the enzyme. To evaluate allosteric inhibition, or differentiate between ATP competitive and non-competitive inhibition, we used an IC_50_ ATP-shift assay (**Fig. 6**). IC_50_ values of the two confirmed selective hits were determined using three ATP concentrations – 10 μM, 100 μM and 500 μM, after optimizing enzyme and substrate concentrations for each ATP concentration. A ∼35 fold reduction in inhibitory potency was observed in both compounds (**1** and **2**) with increasing ATP concentrations. Similarly, a ∼20 fold reduction in inhibitory potency was observed in the reference inhibitor Silmitasertib (CX-4945), known to be an ATP-competitive inhibitor [8, 12]. The potency of putative allosteric inhibitors was expected to remain unchanged with respect to ATP concentration. Hence, this ATP-dependent shift in IC_50_ values of our selective hit compounds suggested that compounds **1** and **2** were ATP-site-directed.

**Figure 6.**
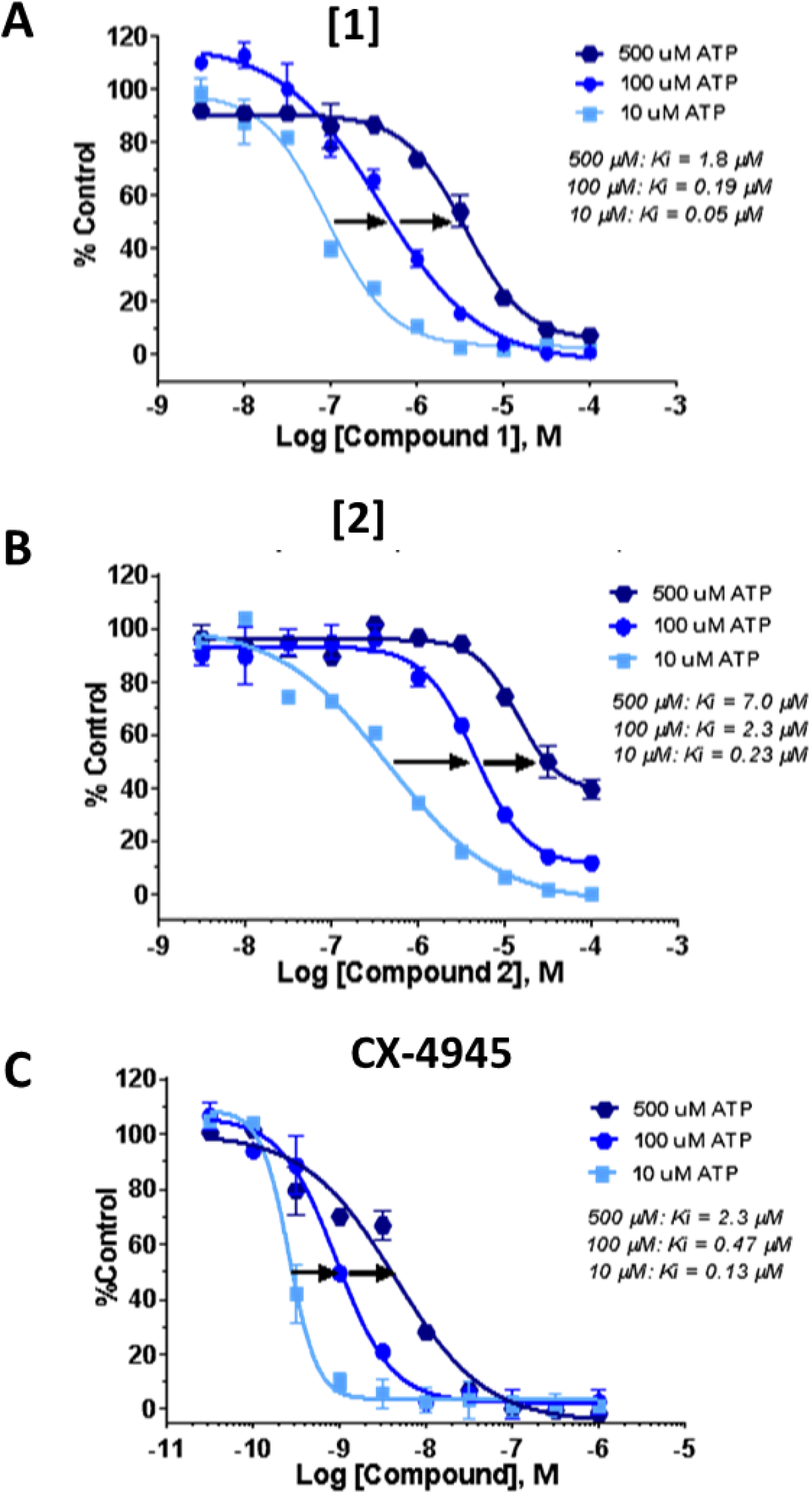
ATP shift assay to distinguish between allosteric and ATP-site HTS hits. IC_50_ values of the two confirmed selective hits were determined using three ATP concentrations – 10 μM, 100 μM and 500 μM. (A,B) A ∼35 fold reduction in inhibitory potency was observed in both compounds (1 and 2) with increasing ATP concentrations. (C) A ∼20-fold reduction in inhibitory potency was observed in the reference inhibitor silmitasertib (CX-4945), a known ATP-competitive inhibitor. This ATP-dependent shift in IC_50_ values of our selective hit compounds suggests that these compounds are ATP-site-directed.

It is not surprising that the allosteric kinase inhibitor-like library screen used in our study turned up ATP competitive inhibitors. This library was based on an unbiased clustering and selection of compounds similar to known allosteric kinase inhibitors. This manner of library construction can lead to a mixed bag of inhibitor types based on the content of the reference library from which they are selected. Though it is available from at least three suppliers, our best compound, **1**, has not previously been reported in the literature (SciFinder^n^, accessed 12-07-2023). Additionally, although some CK2α’ allosteric inhibitors have been previously reported [47–49], a recent structural study has revealed that most of the previously proposed allosteric CK2 inhibitors might indeed act through the ATP site [46]. On the other hand, a recent study has shown that the heterogeneity within conformationally flexible regions of a protein kinase could confer selectivity between highly homologous kinases [58]. Discrete amino acid changes within the ATP site of highly homologous forms of the SFK kinase family have provided isoform selectivity to ATP-competitive inhibitors by allosterically influencing the global conformation of the protein [58]. Therefore, it is possible that ATP-competitive inhibitors could confer allosteric selectivity between CK2α’ and CK2α.

Since the ATP binding site between CK2α’ and CK2α is highly homologous (**Fig. 1B**) we conducted computational induced fit docking with **1** to identify potential binding differences between CK2α’ and CK2α that could explain the enhanced selectivity of this compound for CK2α’. The selectivity of **1** for CK2α’ can be rationalized by key hydrogen bonding interactions between the pyrazine moiety and Y116 that is not present in CK2α (**Fig. 7**).

**Figure 7.**
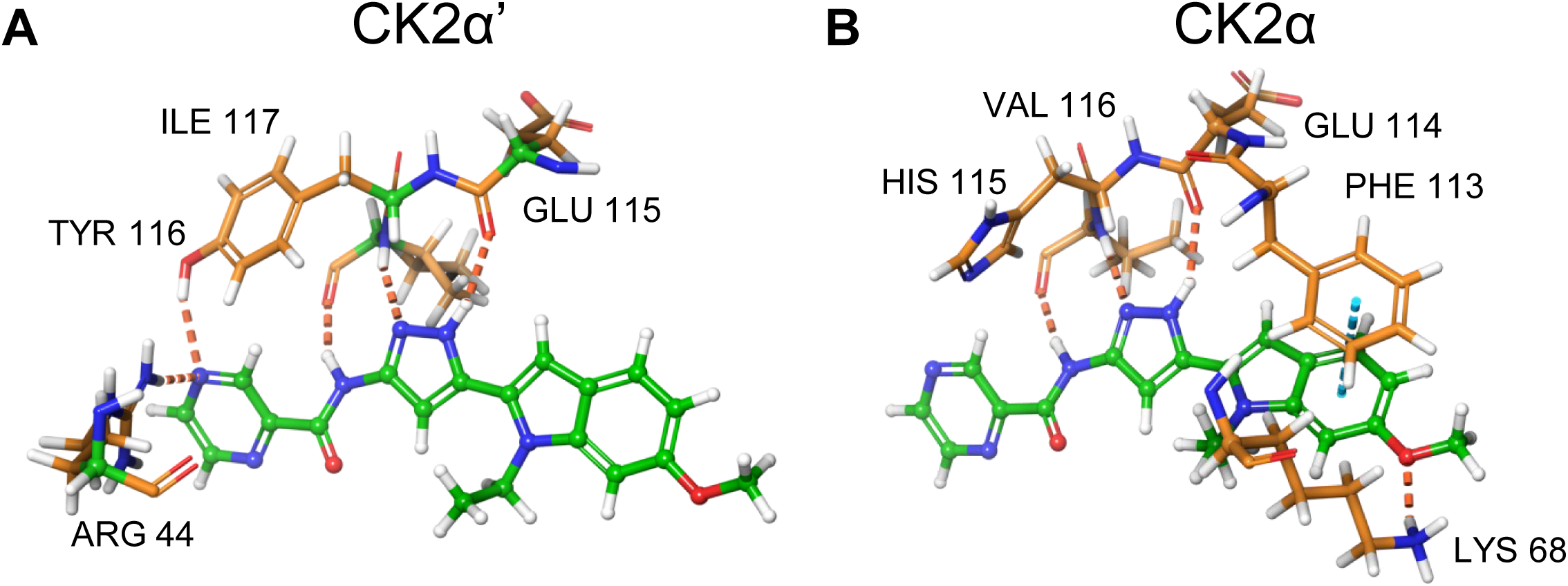
View of the predicted binding mode of **1** with (A) CK2α’, and (B) CK2α that can be used to rationalize the preference of **1** for CK2α’. (Maestro, Schrodinger, 2023-2, Induced Fit Docking, IFD.)

Five different analogs, one for compound **1** (**10**), and four analogs for compound **2** (**11-14**) were purchased and tested to determine if specific structural components were responsible for the selectivity of **1** and **2** towards CK2α’ (**Fig. 8**). All tested analogs lost inhibitory properties on CK2α’, demonstrating the relevance of the pyrazine moiety present in **1** and **2** but not present in the analogs. SAR (IC_50_ CK2α’) of the analogs **10** and **11** can be rationalized in the same fashion as shown for **1** and **2**. Induced fit docking and scoring of these compounds suggests that while both **1** and **2** can form H-bonds with the key Y116, neither **10**, nor **11** can do so (**Fig. 9**). Confirmation of these explanations awaits further analog synthesis and X-ray crystallography. Some potent CK2α’ inhibitors, including CX-4945, feature a carboxylic acid moiety (**Supplementary** Fig. 13) that interacts with K68 (as exemplified in PDB id: 6hme) in the back of the binding pocket[47]. Given that our inhibitors lack such an interaction, it suggests that potency in this series might be increased by including appropriately situated carboxylic acid or carboxylic acid bioisosteres.

**Figure 8.**
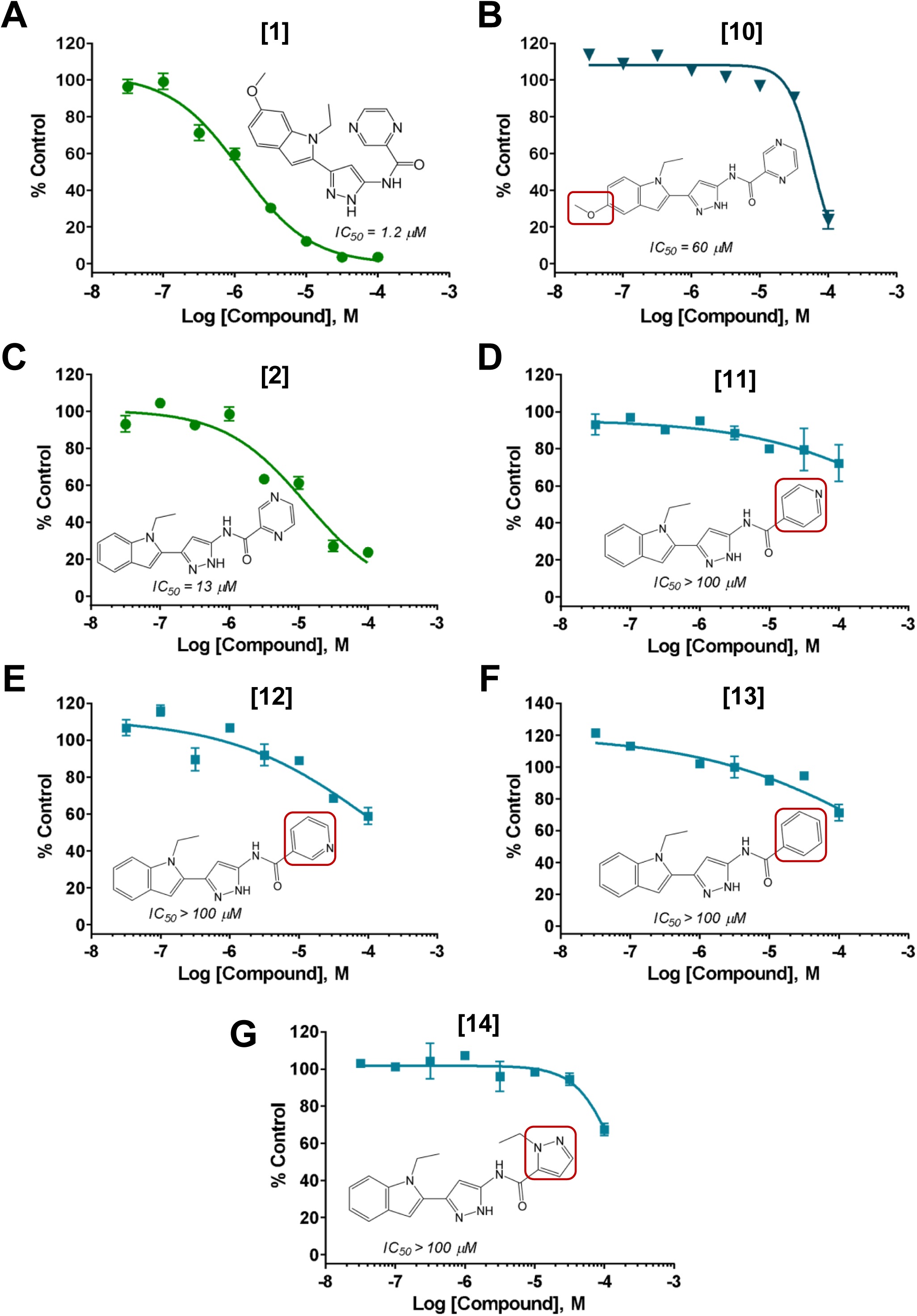
Evaluation of analogs of screening hits to determine SAR. (A,B) **10**, an analog of **1**, showed a 50-fold reduction in inhibitory potency, possibly due to a lack of key hydrogen bonding interactions in the ATP binding site of CK2α‘. (C,D,E,F,G) Similarly, four analogs of **2** also showed significantly reduced potency towards CK2α‘. Red boxes highlight the structural groups modified in the analogs.

**Figure 9.**
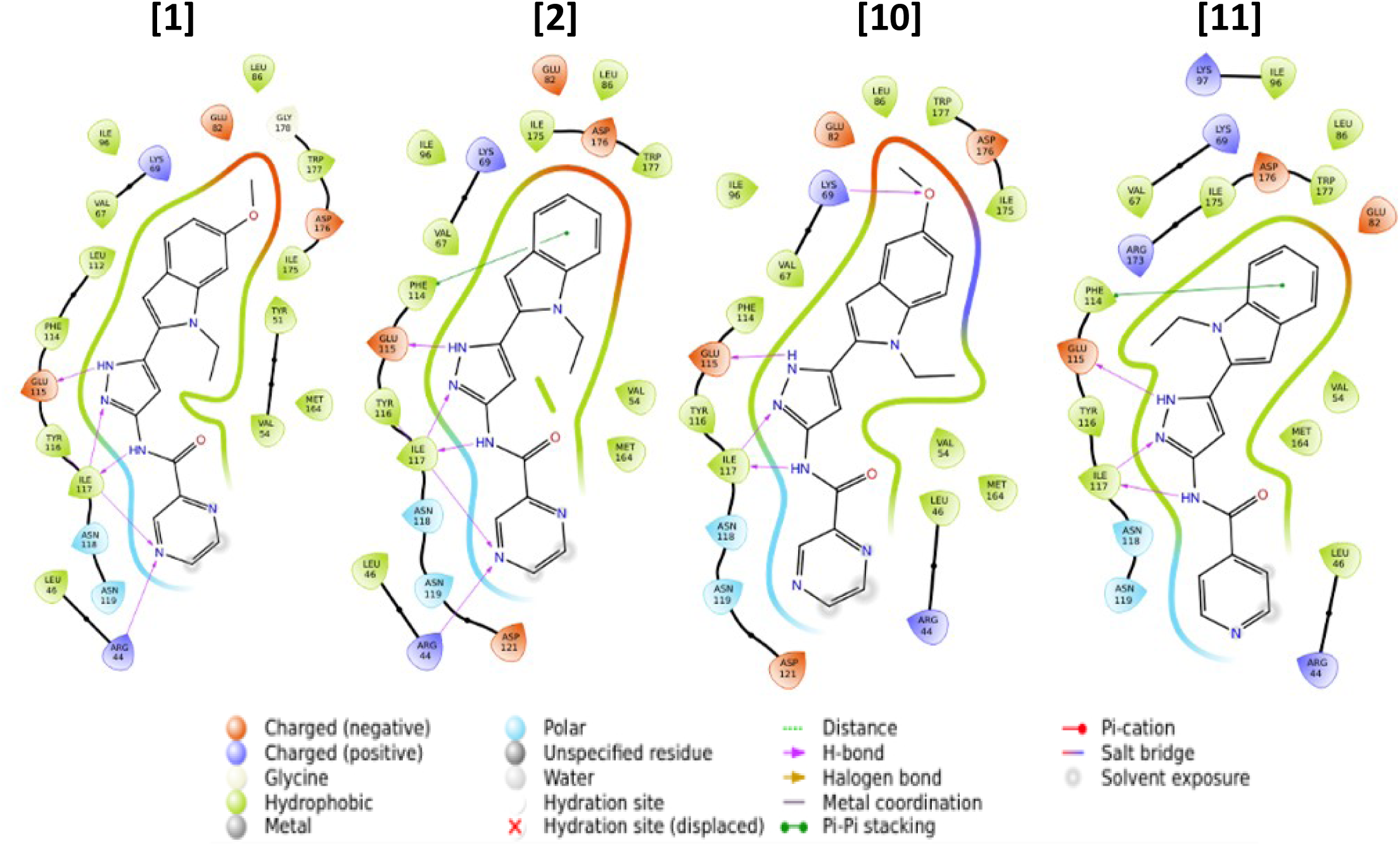
The ligand interaction diagrams computed by Schrodinger, Maestro, for **1**, **2**, **10**, and **11** resulting from IFD (induced fit docking and scoring.)

### Considerations for future assessment of the two CK2**α**’ selective hits

An important factor to be considered in future assessments of **1** and **2**, especially in assessing the inhibition of specific phosphorylated substrates, is the nature of the substrate peptide used in our screening. The commercially available substrate peptide RRRADDSDDDDD has been proven to be a selective substrate for CK2 [53, 59] and has been used in a multitude of studies [60–62]. However, this CK2 consensus sequence is highly promiscuous and is present in more than 20% of the proteome [25, 63]. In addition to the consensus sequence, the recognition mechanism by which kinases identify their substrates depends on 1) the presence of kinase docking motifs outside the catalytic domain that tether enzyme and substrate, and 2) the presence of specific structural determinants surrounding the target site that facilitate enzyme-substrate interaction. These structural determinants are extremely important in the recognition of substrates by Ser/Thr kinases [59, 64]. We previously showed that the HSF1 transcription factor is a substrate for CK2α’. In vitro phosphorylation assays using recombinantly purified CK2α’ and full length HSF1 substrate followed by mass spectrometry analyses revealed several HSF1 residues specifically phosphorylated by CK2α’ including S307, T323 and S333 [30]. Phosphorylation of HSF1 S307 and T323 has previously been shown to have inhibitory effects on HSF1 [65, 66]. We also showed S307 is particularly relevant in the pathological degradation of HSF1 in the context of HD and that CK2α’ genetic knockdown decreased HSF1 S307 phosphorylation and prevented HSF1 degradation [30, 31, 67]. However, we could not detect kinase activity in the ADP-Glo^TM^ assay when using the HSF1 peptide PPQ**S**PRVEEASP containing the S307 residue (bolded). This peptide corresponds to the regulatory domain (RD) of HSF1 [30]. Although the structure of the RD of HSF1 has not been resolved, it is known that this region is highly unstructured due to its dynamic nature [68]. The lack of kinase activity using this peptide could be explained by the unfavorable prolines present in the designed peptide and/or the need for specific structural determinants outside the target site and only present in the fully folded protein. The discrepancies obtained between using fully folded substrates and small synthetic peptide substrates, at least for HSF1, suggests that compound hits obtained using small peptide substrates should be carefully evaluated when assessing inhibitory properties towards specific substrates.

Another factor to be considered is the structural conformation of the CK2 holoenzyme in vivo. CK2 is considered to be a constitutively active enzyme because neither CK2α nor CK2α’ undergo significant conformational changes between active and inactive conformations, as often occurs in other protein kinases [44]. However, it has been proposed that the control of CK2 activity is regulated by a self-inhibitory oligomerization process between the CK2 catalytic subunits and the regulatory subunit CK2β [21, 48]. While monomeric forms of CK2α and CK2α’ are constitutively active, substrate binding can be hampered when they form trimers and/or higher ordered assemblies [69–72]. Oligomerization with CK2β also regulates the substrate specificity of the different catalytic subunits [21, 48, 72]. Since our screening was performed using monomeric forms of CK2α’ it is possible that inhibitory efficacy of compounds 1 and/or 2 over specific substrates might be influenced by the ability of CK2α’ to interact with CK2β in vivo or in vitro. Therefore, future experiments might be directed to elucidate the efficacy of 1 and 2 on the inhibition of specific fully folded CK2α’ substrates and the dependency on the presence of CK2β.

## Materials and Methods

### Kinases and Substrates

Recombinant human GST-CK2α’ (Addgene #27084) and recombinant human GST-CK2α (Addgene #27083) enzymes were produced in-house as previously described [30]. Recombinant Human CKII alpha prime polypeptide His Protein (NBP2-22736) was purchased from Novus Biologicals (Centennial, CO, USA). CK2 Substrate (synthetic peptide RRRADDSDDDDD) was purchased from SignalChem Biotech Inc. (Richmond, BC, Canada). Custom CK2 peptide (RRRDDDSDDD) and HSF1 peptide (PPQSPRVEEASP) were ordered from Biomatik Corporation (Ontario, Canada).

### Chemicals and Assay Components

The ADP-Glo™ Kinase Assay kit (V9104) was purchased from Promega Corporation (Madison, WI, USA). White, 384-well microplates (6005350) were purchased from PerkinElmer Health Sciences Inc. (Shelton, CT, USA). Reference inhibitor CX-4945 (AOB6819) was purchased from AOBIOUS Inc. (Gloucester, MA, USA). 8 repurchased hit compounds (**1-6**, **8** and **9**) were purchased from ChemDiv, Inc. (San Diego, CA, USA). 1 repurchased hit compound (**7**) was purchased from Life Chemicals Inc. (Ontario, Canada). 5 hit analogs (compounds **10-14**) were purchased from ChemDiv, Inc. (San Diego, CA, USA).

### Expression and purification of recombinant proteins

GST-CK2α and GST-CK2α′ expression vectors were purchased from Addgene (pDB1 #27083 and pDB6 #27084, respectively) and recombinant proteins were prepared as previously described [30]. Vectors were transformed into E. coli strain BL21 (DE3). Overnight cultures were diluted 1:100, grown to OD600=0.6 at 37 °C and induced with 1 mM isopropyl 1-thio-β-D-galactopyranoside for 5 h at 37 °C. Cell pellets were lysed in GST equilibration buffer (50 mM Tris, 150 mM sodium chloride, pH 8.0) supplemented, using sonication three times with 30 s bursts. Lysates were cleared by centrifugation at 20,000g for 30 min and incubated with 2 ml (bed volume) of Glutathione agarose beads (Pierce) per liter of culture. Beads were washed twice with equilibration buffer and GST-CK2 subunits eluted with equilibration buffer (50 mM Tris, 150 mM sodium chloride, pH 8.0) supplemented with 10 mM reduced Glutathione. Eluted proteins were concentrated using 5,000LJMWCO centricon, aliquoted (∼ 5mg/ml), flash-frozen in N2 and stored at −80 °C.

### ADP Glo**™** standard curve

ATP to ADP standard curves were prepared in kinase reaction buffer (50 mM Tris-HCl, pH 7.5, 10 mM MgCl2, 0.1 mM EDTA, 100 mM NaCl, 2 mM DTT, 0.01% Triton X-100) to assess the linearity of the assay, and to estimate the amount of ADP produced in the kinase reaction. The standard samples used to generate an ATP-to-ADP conversion curve were created by combining the appropriate volumes of ATP and ADP stock solutions (12 standards ranging from 100% ADP + 0% ATP to 0% ADP + 100% ATP). 5 μL of each standard was added to the assay plate in triplicate using a manual pipette. Next, 5 ul of the ADP Glo™ reagent was added, followed by incubation at RT for 40 mins. 10 ul of the kinase detection reagent was then added, followed by a 60-minute incubation at RT. The resulting luminescence was measured on an EnSpire multimode plate reader (PerkinElmer, Waltham, MA). The data was analyzed and plotted in Prism (Graphpad Software, MA, USA) using linear regression analysis.

### ADP Glo kinase activity assay

All kinase reactions were performed in a 5 ul assay volume at RT. 2.5 µL of 2X recombinant human GST-CK2α’ enzyme was first added to the assay wells using a manual pipette, and the reaction was initiated by the addition of 2X CK2 peptide substrate + ATP. The plate was then centrifuged at 1000 rpm for 30 s (5804/ 5804 R – Benchtop Centrifuge, Eppendorf North America, CT, USA), mixed on a plate shaker (Siemens DPC MicroMix 5) for 1 minute at medium speed, and incubated at RT for 30 min. An ATP to ADP standard curve was included on the assay plate. At the end of the incubation, the reaction was stopped by adding 5 µL of ADP-Glo™ reagent, and the plate was centrifuged again at 1000 rpm for 30 s, mixed on the plate shaker for 1 minute at medium speed, and incubated at RT for 40 min. 10 µL of ADP-Glo™ kinase detection reagent was then added, and the plate was covered and incubated at RT for 1 h. The resulting luminescence was measured on the EnSpire multimode plate reader, and the data was analyzed and plotted in Prism using interpolation of the standard curve and linear regression analysis.

### Determination of standard conditions for the HTS assay

Determination of optimal enzyme concentration was carried out using 80 µM of CK2 peptide substrate and 200 μM ATP with increasing concentration of recombinant human GST-CK2α’ enzyme (10 nM to 300 nM). The reaction was allowed to proceed at RT for 30 minutes. The ADP Glo™ and kinase detection reagents were then added according to the manufacturer’s instructions, and the resulting luminescence was measured on an EnSpire multimode plate reader. The % conversion was calculated in MS Excel, and the data was plotted in Prism using linear regression analysis and interpolation of the standard curve.

Determination of the Michaelis–Menten constant (K_M_) of CK2 peptide substrate was carried out using 40 nM recombinant human GST-CK2α’ enzyme and 200 μM ATP with increasing concentrations of CK2 peptide substrate (0 – 800 μΜ). The reaction was allowed to proceed at RT for 30 minutes. The ADP Glo™ and kinase detection reagents were then added according to the manufacturer’s instructions, and the resulting luminescence was measured on an EnSpire multimode plate reader. The % conversion was calculated in MS Excel, and the K_M_ was calculated by fitting the data to the classic Michaelis–Menten equation in Prism.

Determination of the Michaelis–Menten constant (K_M_) of ATP was carried out using 20 nM recombinant human GST-CK2α’ enzyme and a saturating concentration (600 μM) of CK2 peptide substrate, with increasing concentrations of ATP (7.8 μΜ – 1 mM). The reaction was allowed to proceed at RT for 30minutes. The ADP Glo™ and kinase detection reagents were then added according to the manufacturer’s instructions, and the resulting luminescence was measured on an EnSpire multimode plate reader. The % conversion was calculated in MS Excel, and the K_M_ was calculated by fitting the data to the classic Michaelis–Menten equation in Prism.

### HTS assay validation

The controls used for the standard HTS assay were a high signal (no inhibitor) negative control, a low signal (no enzyme) background control, and a mid-signal (IC50 concentration of reference inhibitor CX-4945) positive control. The vertical validation assay plate comprised of alternating columns of the high, low, and mid controls dispensed in identical sets across the plate using the Echo 550 dispenser (Beckman Coulter, Brea, CA) for compounds/DMSO, and the Multidrop Combi nL (Thermo Fisher Scientific, Waltham, MA) liquid dispenser for assay reagents. CX-4945 was added to the positive control wells at a final concentration of 5 nM, and an equivalent volume of DMSO was added to the high and low control wells for a final DMSO concentration of 3.2%. 2.5 μL of kinase assay buffer was added to all low control wells, and 2.5 μL of recombinant human GST – CK2α’ enzyme at a final concentration of 40 nM was added to the high and mid control wells using a Multidrop Combi nL liquid dispenser. The plate was centrifuged at 1000 rpm for 30 s, mixed on the plate shaker for 1 minute at medium speed, and incubated at RT for 60 min. The kinase reaction was then initiated by the addition of CK2 peptide substrate and ATP at a final concentration of 80 µM and 200 µM respectively. The reaction was allowed to proceed at RT for 30 minutes. The ADP Glo™ reagent and kinase detection reagents were then added according to the manufacturer’s instructions, and the resulting luminescence was measured on an EnSpire multimode plate reader. The Z′ factor was calculated using the following equation:

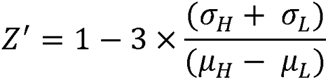

where *σ*_H_ and *σ*_L_ are the standard deviations of the high and low controls, respectively, and *µ*_H_ and *µ*_L_ are their respective means.

A pilot screen was performed with the TOCRIS kinase inhibitor toolbox (80 compounds; (Cat# 3514, TOCRIS Bioscience, Minneapolis, MN) and the GSK kinase inhibitor PKIS2 collection (473 compounds; GlxoSmithKline, Research Triangle Park, NC). Compounds were added to 384-well microplates using the Echo 550 dispenser at a final concentration of 10 µM. High, low and mid signal controls were included on all pilot screen plates. The assay was performed using standard HTS assay conditions as described above, and repeated 3X on three independent days. Z’ values and the hit rate were calculated for each independent run.

### High throughput screen

The optimized ADP Glo™ standard activity assay was used to screen the Allosteric Kinase Inhibitor Collection (ChemDiv, Inc., San Diego, CA, USA) of 28,812 compounds for allosteric inhibitors. The reference inhibitor CX-4945 was included as a positive control on every screening plate in addition to the high and low controls. For the HTS, compounds were added to 384-well microplates using the Echo 550 dispenser at a final concentration of 10 µM. A Multidrop Combi nL (Thermo Fisher Scientific, Waltham, MA) liquid dispenser was used to add 2.5 μL of recombinant human glutathione S-transferase (GST) – CK2α’ enzyme diluted in kinase assay buffer at a final concentration of 40 nM, and pre-incubated for 60 min at RT after shaking for 1 minute at medium speed on a plate shaker. The kinase reaction was then initiated by the addition of 2.5 μL of CK2 peptide substrate at a final concentration of 80 µM in kinase assay buffer containing 200 μM ATP, and incubated for 30 minutes at RT. The ADP Glo™ and kinase detection reagents were then added according to the manufacturer’s instructions, and luminescence was measured on the EnSpire multimode plate reader. %Inhibition was calculated using the following equation:

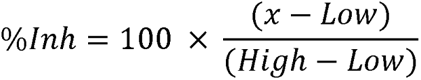

where x is the measured luminescence in the compound well, Low is the average signal from the no-enzyme control column, and High is the average signal from the no-inhibitor control column.

Compounds producing greater than 50% inhibition were identified as screening hits.

### Dose response assays for hit validation

To confirm and determine inhibitory potency of screening hits, compounds were cherry-picked from DMSO stocks and added to assay plates using the Echo 550 dispenser at eight concentrations in duplicate (100 μM – 30 nM). The reference inhibitor CX-4945 was also added as a positive control using the Echo 550 dispenser at eight concentrations in duplicate (1 μM – 0.3 nM) on every assay plate. The final DMSO concentration was 3.5%, and the assay was performed at standard HTS assay conditions. IC_50_ values were calculated using a four-parameter nonlinear regression analysis in Prism.

### Luciferase reaction interference assay

Compounds that interfere with the luciferase reaction were detected by assaying a duplicate set of assay wells without the enzyme, and using ADP as the substrate. Prioritized confirmed hits were dispensed in triplicate using the Echo 550 dispenser at a final concentration of 20 µM into two assay plates as duplicate sets of assays. One plate was tested with recombinant human glutathione S-transferase (GST) – CK2α’ enzyme and CK2 peptide substrate at standard HTS assay conditions, while the second plate was tested with a fixed concentration (20 µM) of ADP. High and low (no-enzyme/ADP) controls were included on both plates. Percent inhibition was calculated for compounds on both plates using the equation described above. Compounds that produced apparent inhibition in the absence of the enzyme when ADP was used as a substrate were identified as interfering compounds.

### Selectivity assay

Prioritized, confirmed screening hits that did not interfere with the luciferase reaction, either cherry-picked from compound library DMSO stocks, or fresh DMSO stocks of repurchased powders were added to assay plates using the Echo 550 dispenser at eight concentrations in duplicate (100 μM – 30 nM). Two identical assay plates were created for each set of 20 compounds. The reference inhibitor CX-4945 was also added as a positive control using the Echo 550 dispenser at eight concentrations in duplicate (1 μM – 0.3 nM) on every assay plate. For each set of compounds, one plate was tested with recombinant human glutathione S-transferase (GST) – CK2α’ enzyme, while the other was tested with recombinant human glutathione S-transferase (GST) – CK2α enzyme under standard assay conditions. IC_50_ values were calculated using a four-parameter nonlinear regression analysis in Prism. Compounds that produced greater than a 3-fold difference in potency, or 2-fold difference in efficacy were identified as selective compounds.

### Mode of action assay: IC_50_ shift

To evaluate the mode of action of inhibition of selective compounds, three identical 12-point dose responses (100 μM to 0.3 nM) were tested in the presence of 10 μM ATP, 100 μM ATP, and 500 μM ATP. Optimal enzyme concentration for each ATP condition was determined as described above, and standard assay conditions were used for the rest of the assay parameters. IC_50_ values were calculated using a four-parameter nonlinear regression analysis in Prism. Compounds that displayed a shift in IC_50_ values dependent on the ATP concentration were identified as ATP-competitive.

### NMR

Experiments were performed on a 400/100 MHz instrument. NMR spectra were processed with the MestReNova software. Chemical shifts are shown in ppm relative to DMSO-d^6^ (2.50 ppm for ^1^H NMR).

### Purity Analysis

The UPLC analyses and mass spectra (LC-MS) were obtained on Waters ACQUITY system (Waters, Milford, CT,USA) with a QDa Mass Detector with ESI inlet and UV PDA detector. The Waters UPLC BEH C18, 1.7µm (2.1 x 50 mm), column was used at 40 °C temperature. The samples were dissolved in between 200 µL and 600 µL of DMSO depending on the solubility of the sample. The sample was then diluted to approximately 0.1 mg/mL in MeOH for injection onto the LCMS. The LCMS parameters were as follows: mobile Phase A (10 mM ammonium bicarbonate in water), mobile Phase B (ACN or MeOH), a flow rate of 0.6 mL/min, injection volume 7.5 µL, run time 6.0 min. Gradient Operation as follows: Hold at 95:5 A:B for 0.5 min. Linear gradient to 5:95 A:B for 3 min. Hold at 5:95 A:B for 0.5 min. Linear gradient to 95:5 A:B for 0.5 min. Hold at 95:5 A:B for 1.5 min. The mass detector was run at a positive scan from 150 – 1000 Da. The PDA quantified the peaks at 254 nm.

### Computational Studies

All computational studies were performed using Schrödinger 2022-2 or 2023-2 and the utilities therein. The reported crystal structures of CK2α (PDBid: 6hme) and CK2α’ (PDBid: 6hmc) were used as starting points for docking and scoring studies. (1) These proteins were prepared using the default parameters in the Protein Preparation wizard in Maestro. This preparation scheme led to the holo, dry (no waters incorporated) structures used for all subsequent studies. Induced fit dockings (IFD)[73] were performed using the default parameters in Maestro (extended sampling, which generates up to 80 poses using automatic docking settings, was employed). The IFD structures with the lowest docking energies were selected to analyze key protein-ligand interactions.

### Safety

The authors are not aware of significant hazards or risks associated with the reported work.

### Abbreviations

CK2α; CK2 alpha, CK2α’; CK2 alpha prime, IFD; induced fit docking, GST: glutathione S-transferase, HD; Huntington’s disease, IC_50_; half-maximal inhibitory concentration, HTS: high throughput screening.

## Author Information

### Corresponding author

*Rocio Gomez-Pastor*; University of Minnesota, School of Medicine, Department of Neuroscience. Minneapolis, 55455, Minnesota, USA.

### Authors

*Deepti Mudaliar*; Department of Medicinal Chemistry, Institute for Therapeutics Discovery and Development, University of Minnesota, Minneapolis, Minnesota 55414, United States.

*Rachel Mansky*; University of Minnesota, School of Medicine, Department of Neuroscience. Minneapolis, 55455, Minnesota, USA.

*Angel White*; University of Minnesota, School of Medicine, Department of Neuroscience. Minneapolis, 55455, Minnesota, USA.

*Grace Baudhuin*; University of Minnesota, School of Medicine, Department of Neuroscience. Minneapolis, 55455, Minnesota, USA.

*Jon Hawkinson*; Department of Medicinal Chemistry, Institute for Therapeutics Discovery and Development, University of Minnesota, Minneapolis, Minnesota 55414, United States.

*Henry Wong*; Department of Medicinal Chemistry, Institute for Therapeutics Discovery and Development, University of Minnesota, Minneapolis, Minnesota 55414, United States.

*Michael A. Walters*; Department of Medicinal Chemistry, Institute for Therapeutics Discovery and Development, University of Minnesota, Minneapolis, Minnesota 55414, United States.

### Author Contributions

Rocio Gomez-Pastor and Michael Walters designed the experiments, obtained funding, and provided guidance in data interpretation. Michael Walters performed induced fit docking analyses and provided guidance in medicinal chemistry. Jon Hawkinson and Henry Wong provided guidance in compound screening and data interpretation. Deepti Mudaliar performed the HTS ADP-Glo™ screening and hits validation. Rachel Mansky produced the recombinant CK2α and CK2α’ proteins. Angel White and Grace Baudhuin participated in the validation of compounds 1 and 2.

## Acknowledgments

The authors acknowledge the Minnesota Supercomputing Institute (MSI) at the University of Minnesota (http://www.msi.umn.edu) for providing resources that contributed to the research results reported within this paper. We also thank Andrew Goode for performing purity analyses on repurchased compounds and Dr. Narsihmulu Cheryala for performing NMR analyses on compounds 1 and 2. We thank the NIH (R21NS116260) for funding.

