## Supplementary Table 1, Supplementary figures 1-13 for "Identification of CK2α’ selective inhibitors by the screening of an allosteric-kinase-inhibitor-like compound library"

### Supplementary Material

**Supplementary Table 1.** Interference assay to detect compounds that interfered with the ADP-Glo™ luciferase reaction. None of the tested compounds produced any significant inhibition in the absence of enzyme.

| Compound ID | % Inhibition | |
|---|---|---|
| | + Enzyme | +ADP |
| GPHR-00147239 | 90.8 | 5.2 |
| GPHR-00333025 | 84.5 | 2.7 |
| GPHR-00333745 | 99.1 | 1.6 |
| GPHR-00333753 | 88.2 | -1.0 |
| GPHR-00334441 | 91.2 | -1.0 |
| GPHR-00336048 | 96.1 | -3.7 |
| GPHR-00336409 | 99.6 | -1.2 |
| GPHR-00337058 | 98.2 | -4.4 |
| GPHR-00337062 | 96.4 | -4.0 |
| GPHR-00337416 | 96.3 | -6.2 |
| GPHR-00337388 | 84.4 | -5.2 |
| GPHR-00337420 | 98.3 | -8.6 |
| GPHR-00337422 | 91.3 | -6.4 |
| GPHR-00337392 | 97.7 | -5.9 |
| GPHR-00337394 | 91.0 | -4.3 |
| GPHR-00337426 | 98.5 | -8.5 |
| GPHR-00337447 | 98.6 | 4.1 |
| GPHR-00337530 | 96.3 | 0.7 |
| GPHR-00337451 | 89.7 | 1.1 |
| GPHR-00337483 | 74.4 | -2.1 |
| GPHR-00337662 | 84.7 | -1.7 |
| GPHR-00337535 | 97.1 | -4.3 |
| GPHR-00337489 | 92.3 | -3.1 |
| GPHR-00339923 | 97.2 | -4.2 |
| GPHR-00341143 | 81.3 | -4.9 |
| GPHR-00341145 | 92.2 | -7.6 |
| GPHR-00341121 | 78.0 | -6.6 |
| GPHR-00342435 | 90.4 | -7.8 |
| GPHR-00342230 | 69.9 | -7.2 |
| GPHR-00342300 | 73.6 | -6.9 |
| GPHR-00343938 | 90.5 | -6.1 |
| GPHR-00343919 | 74.7 | -9.2 |
| GPHR-00344034 | 71.5 | 5.4 |

| Compound ID | % Inhibition | |
|---|---|---|
| | + Enzyme | +ADP |
| GPHR-00344195 | 94.3 | 0.4 |
| GPHR-00344201 | 83.8 | 1.0 |
| GPHR-00327079 | 99.2 | -2.3 |
| GPHR-00344426 | 84.7 | -0.9 |
| GPHR-00344428 | 98.5 | -6.4 |
| GPHR-00344444 | 64.5 | -2.9 |
| GPHR-00344589 | 91.3 | -5.4 |
| GPHR-00344450 | 78.3 | -5.4 |
| GPHR-00344570 | 96.2 | -8.6 |
| GPHR-00344468 | 89.7 | -7.0 |
| GPHR-00344572 | 84.5 | -9.1 |
| GPHR-00344892 | 98.8 | -8.1 |
| GPHR-00344912 | 92.6 | -7.9 |
| GPHR-00344810 | 92.5 | -7.1 |
| GPHR-00344918 | 95.4 | -10.5 |
| GPHR-00345919 | 80.5 | 7.2 |
| GPHR-00347991 | 86.0 | 2.9 |
| GPHR-00347755 | 81.4 | 3.2 |
| GPHR-00347963 | 73.8 | 1.0 |
| GPHR-00352937 | 93.3 | -1.7 |
| GPHR-00012891 | 97.5 | -4.0 |
| GPHR-00037954 | 65.3 | -1.5 |
| GPHR-00353237 | 93.5 | -2.3 |
| GPHR-00353219 | 95.3 | -4.8 |
| GPHR-00353287 | 75.6 | -5.7 |
| GPHR-00353494 | 97.9 | -5.5 |
| GPHR-00353528 | 86.9 | -7.3 |
| GPHR-00328352 | 79.7 | -7.0 |
| GPHR-00353524 | 79.2 | -6.6 |
| GPHR-00353752 | 56.2 | -5.4 |
| GPHR-00353812 | 80.1 | -7.9 |
| GPHR-00353775 | 78.4 | 8.3 |
| CX-4945 | 98.7 | 4.1 |

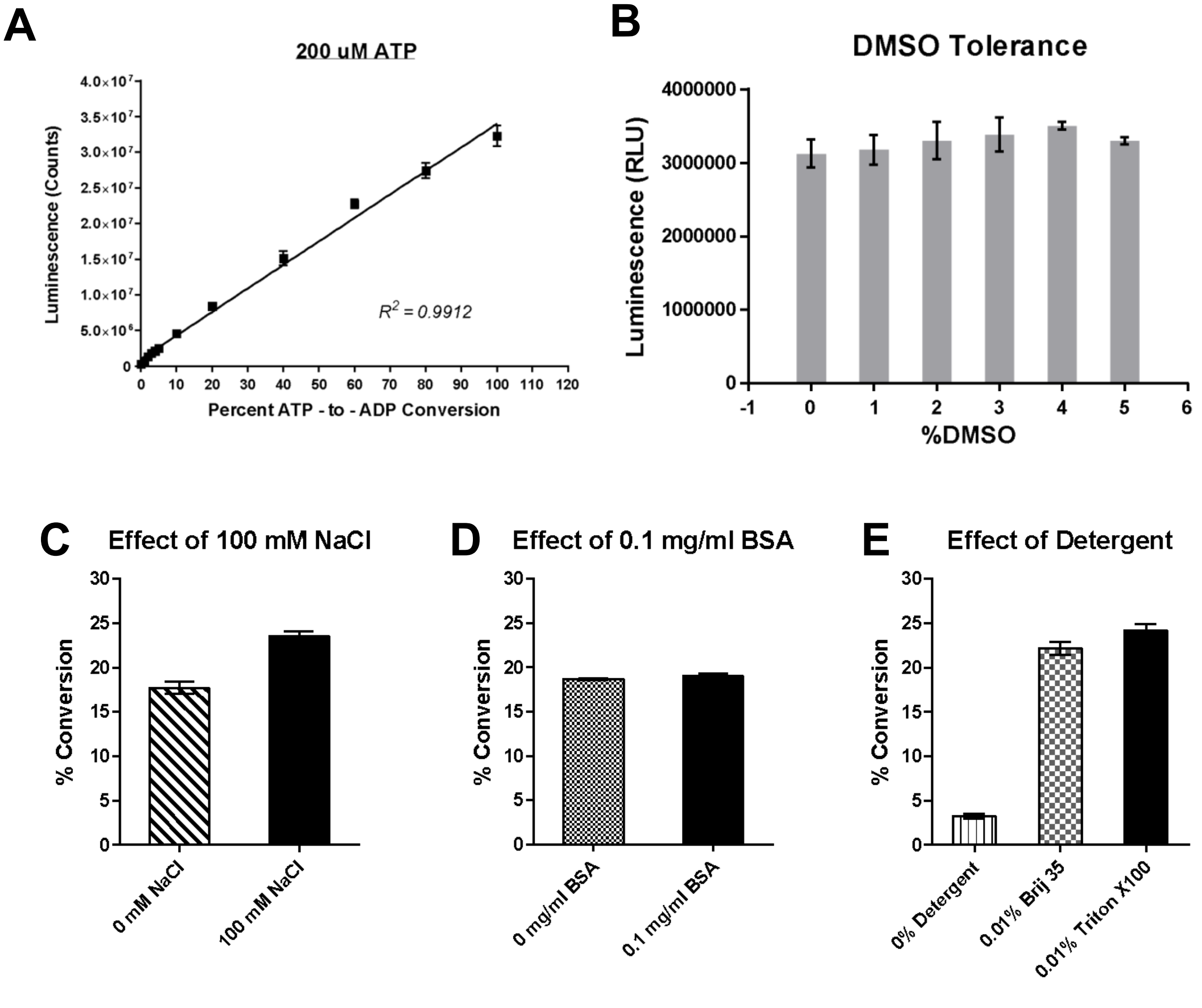


**Supplementary Figure 1. CK2a'-dependent ADP-Glo assay optimization.** (A) The ATP–ADP standard curve used to calculate the amount of ADP produced in the kinase enzyme reaction. The amount of luminescence (measured as RLU) corresponds to the conversion of ATP to ADP based on the ATP concentration used in the reaction. (B) DMSO tolerance assay. The assay showed no detrimental effect of DMSO for up to 5%. (C) Buffer optimization. The addition of 100 mM NaCl and 0.01% detergent resulted in a 1.5 to 5-fold increase in percent conversion, while the addition of BSA did not affect the enzyme activity.

[1]

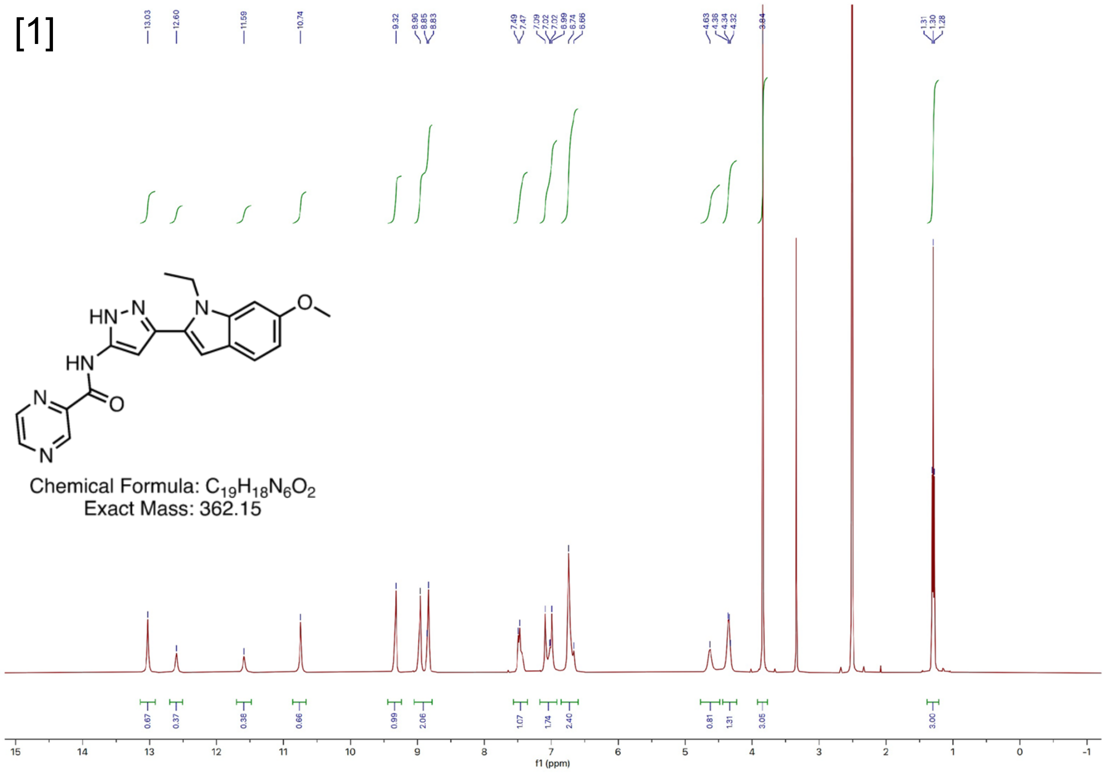


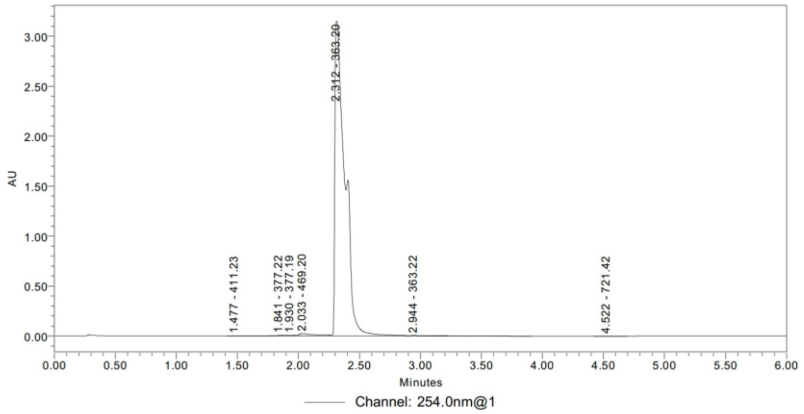


| | RT | Area | % Area | Height | Base Peak (m/z) |
|---|---|---|---|---|---|
| 1 | 1.477 | 4644 | 0.02 | 1249 | 411.23 |
| 2 | 1.841 | 39618 | 0.21 | 7161 | 377.22 |
| 3 | 1.930 | 42320 | 0.23 | 8685 | 377.19 |
| 4 | 2.033 | 247102 | 1.33 | 23423 | 469.20 |
| 5 | 2.312 | 18149416 | 97.37 | 3151176 | 363.20 |
| 6 | 2.944 | 153021 | 0.82 | 7597 | 363.22 |
| 7 | 4.522 | 3160 | 0.02 | 399 | 721.42 |

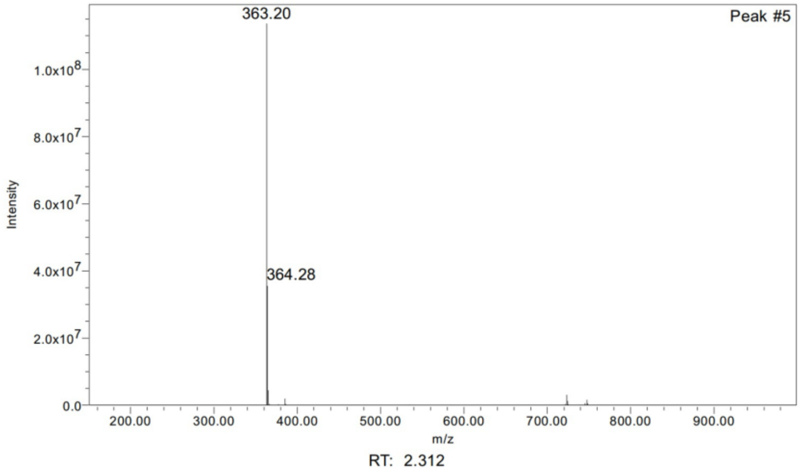


**Supplementary Figure 2. NMR spectra and LC-MS for repurchased compound 1.**

[2]

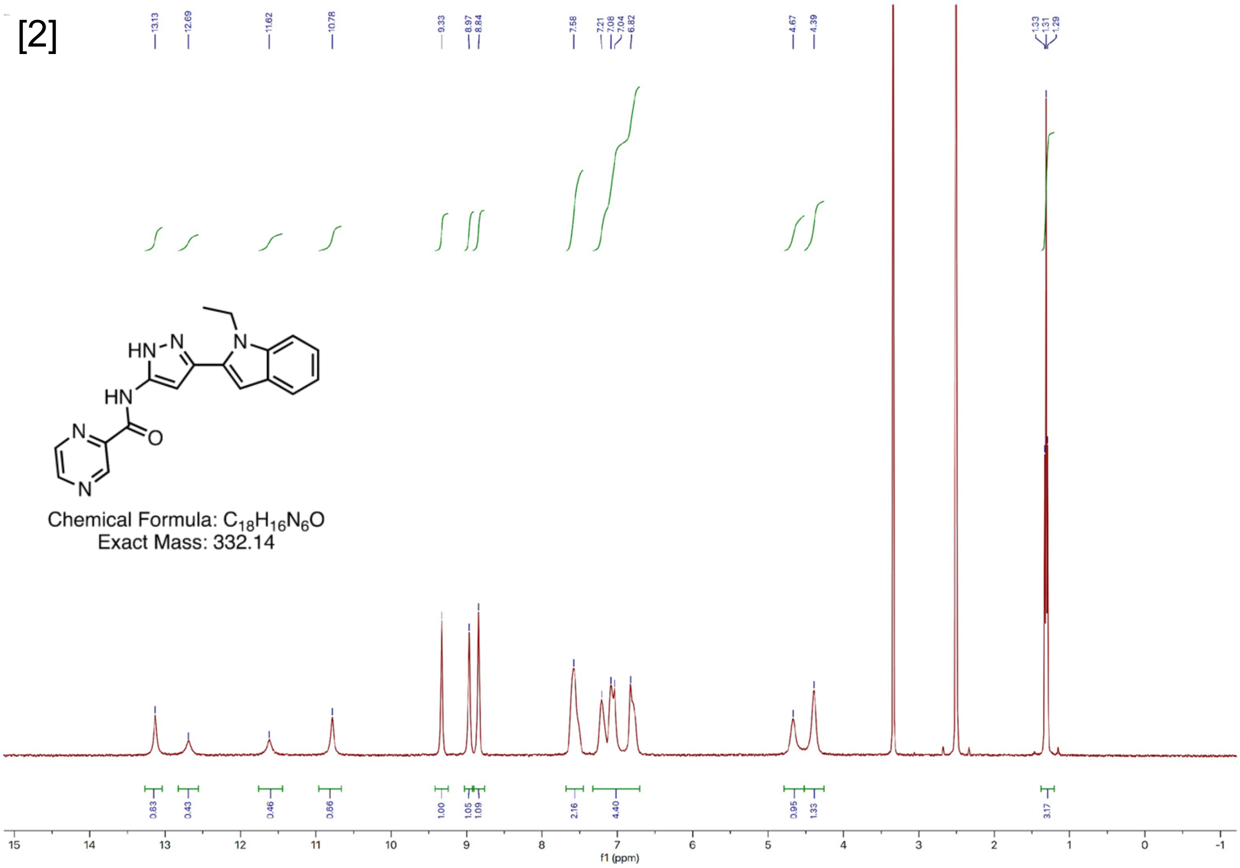


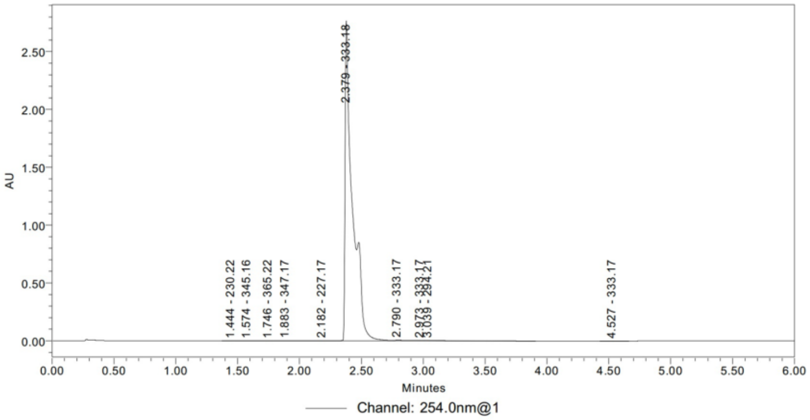


| | RT | Area | % Area | Height | Base Peak (m/z) |
|---|---|---|---|---|---|
| 1 | 1.444 | 1279 | 0.01 | 304 | 230.22 |
| 2 | 1.574 | 1574 | 0.01 | 292 | 345.16 |
| 3 | 1.746 | 3456 | 0.03 | 494 | 365.22 |
| 4 | 1.883 | 12460 | 0.11 | 1549 | 347.17 |
| 5 | 2.182 | 21065 | 0.18 | 1449 | 227.17 |
| 6 | 2.379 | 11537460 | 98.30 | 2769224 | 333.18 |
| 7 | 2.790 | 65131 | 0.55 | 7866 | 333.17 |
| 8 | 2.973 | 8220 | 0.07 | 3960 | 333.17 |
| 9 | 3.039 | 84161 | 0.72 | 4537 | 294.21 |
| 10 | 4.527 | 1654 | 0.01 | 222 | 333.17 |

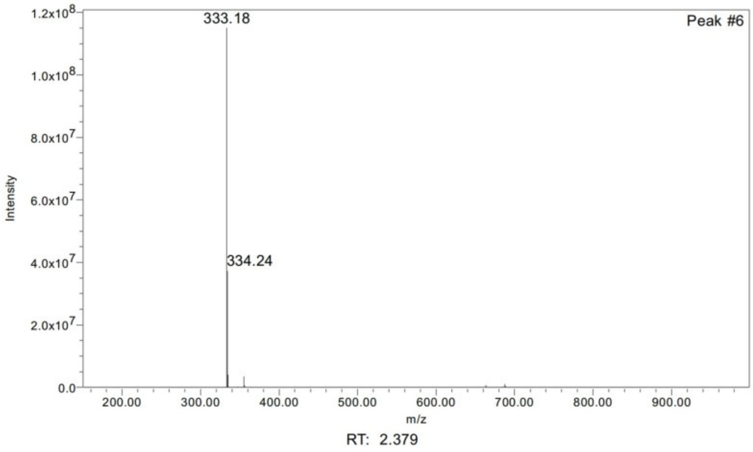


**Supplementary Figure 3. NMR spectra and LC-MS for repurchased compound 2.**

[3]

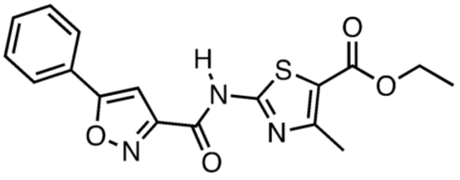

Chemical Formula: $C_{17}H_{15}N_3O_4S$
Exact Mass: 357.08

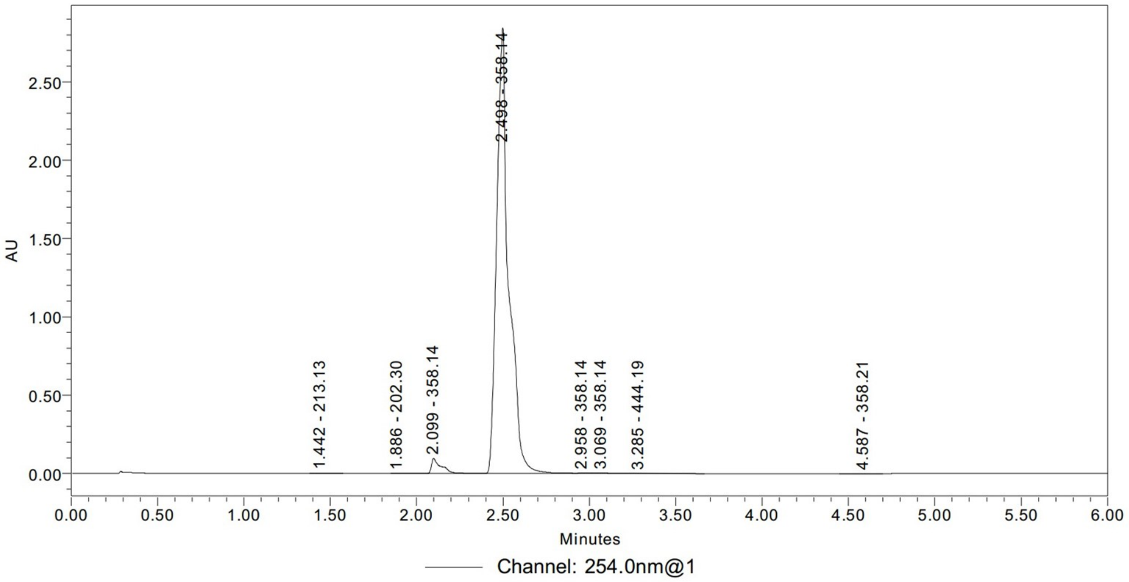


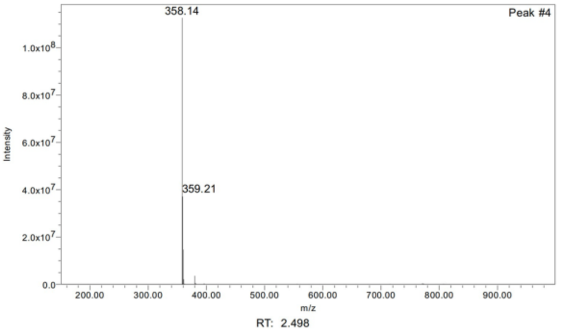


| | RT | Area | % Area | Height | Base Peak (m/z) |
|---|---|---|---|---|---|
| 1 | 1.442 | 4330 | 0.03 | 1150 | 213.13 |
| 2 | 1.886 | 3520 | 0.02 | 706 | 202.30 |
| 3 | 2.099 | 379656 | 2.65 | 97170 | 358.14 |
| 4 | 2.498 | 13856935 | 96.87 | 2845102 | 358.14 |
| 5 | 2.958 | 14142 | 0.10 | 2783 | 358.14 |
| 6 | 3.069 | 29948 | 0.21 | 3077 | 358.14 |
| 7 | 3.285 | 14817 | 0.10 | 1582 | 444.19 |
| 8 | 4.587 | 1486 | 0.01 | 177 | 358.21 |

**Supplementary Figure 4. LC-MS data for repurchased compound 3.**

[4]

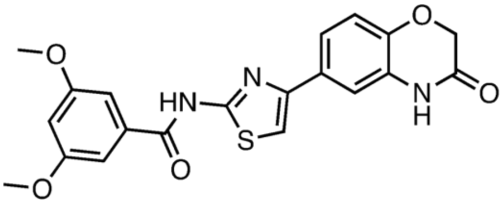


Chemical Formula: $C_{20}H_{17}N_{3}O_{5}S$
Exact Mass: 411.09

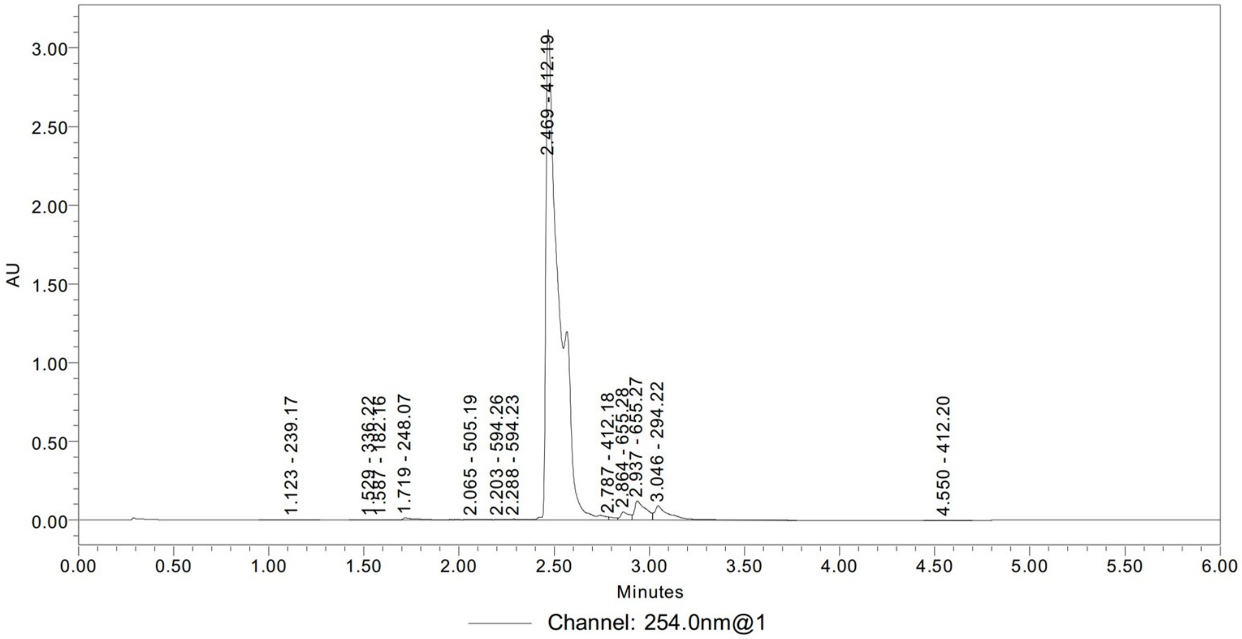


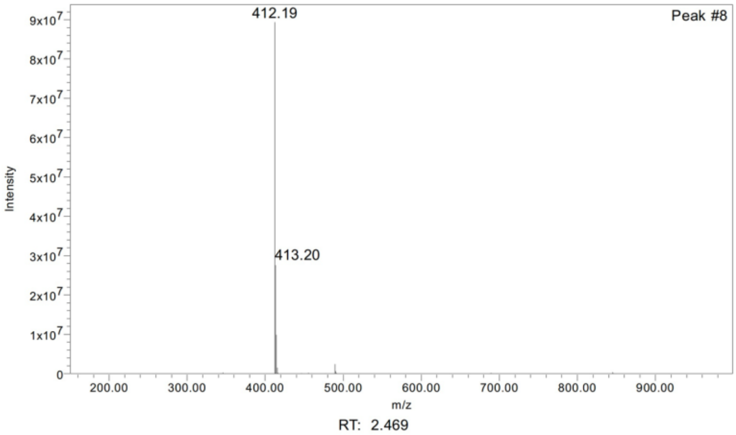


| | RT | Area | % Area | Height | Base Peak (m/z) |
|---|---|---|---|---|---|
| 1 | 1.123 | 5796 | 0.03 | 726 | 239.17 |
| 2 | 1.529 | 1655 | 0.01 | 405 | 336.22 |
| 3 | 1.587 | 3072 | 0.02 | 721 | 182.16 |
| 4 | 1.719 | 70883 | 0.43 | 13537 | 248.07 |
| 5 | 2.065 | 45562 | 0.27 | 6032 | 505.19 |
| 6 | 2.203 | 15411 | 0.09 | 5942 | 594.26 |
| 7 | 2.288 | 46103 | 0.28 | 6641 | 594.23 |
| 8 | 2.469 | 15237190 | 91.53 | 3117707 | 412.19 |
| 9 | 2.787 | 45872 | 0.28 | 19727 | 412.18 |
| 10 | 2.864 | 163893 | 0.98 | 52647 | 655.28 |
| 11 | 2.937 | 491228 | 2.95 | 122422 | 655.27 |

**Supplementary Figure 5. LC-MS data for repurchased compound 4.**

[5]

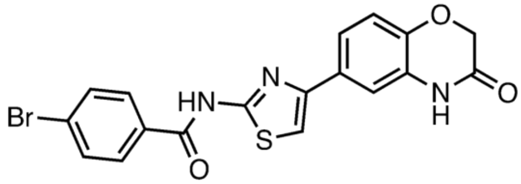


Chemical Formula: $C_{18}H_{12}BrN_3O_3S$
Exact Mass: 428.98

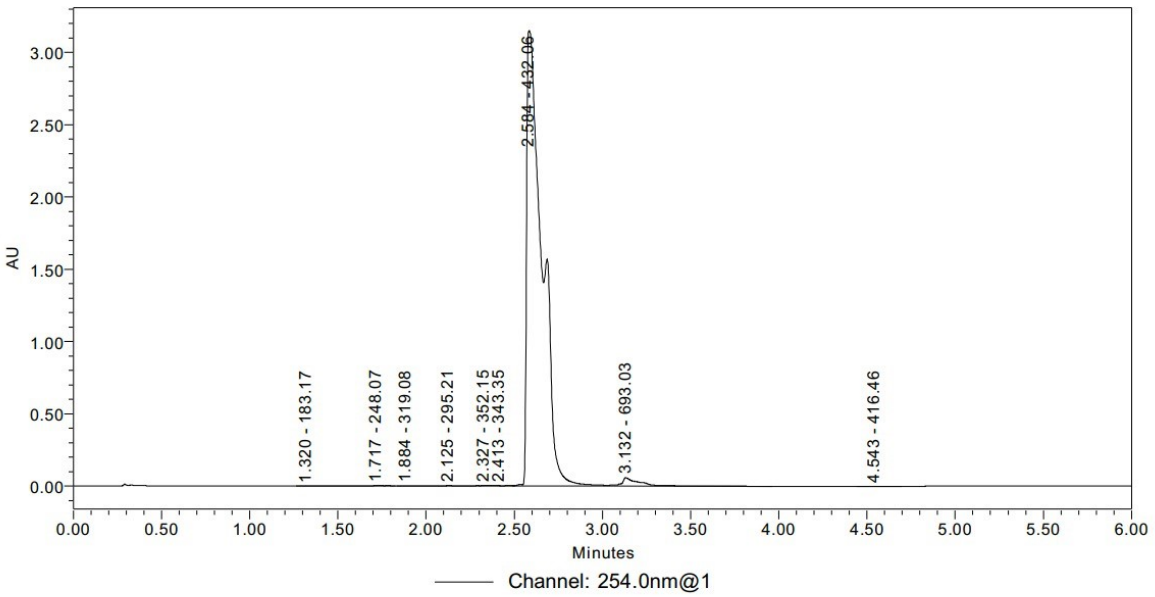


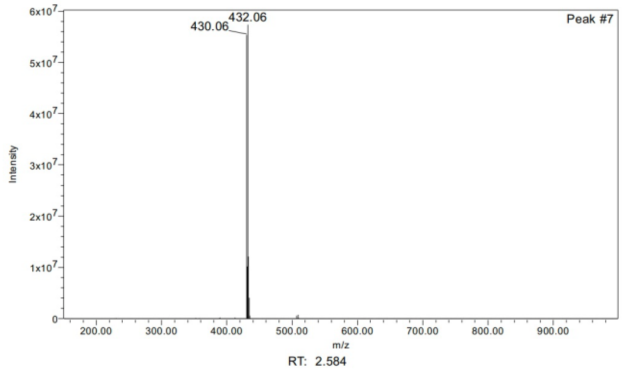


| | RT | Area | % Area | Height | Base Peak (m/z) |
|---|---|---|---|---|---|
| 1 | 1.320 | 1562 | 0.01 | 342 | 183.17 |
| 2 | 1.717 | 24299 | 0.12 | 4450 | 248.07 |
| 3 | 1.884 | 12892 | 0.07 | 1784 | 319.08 |
| 4 | 2.125 | 19438 | 0.10 | 3783 | 295.21 |
| 5 | 2.327 | 33111 | 0.17 | 4853 | 352.15 |
| 6 | 2.413 | 4628 | 0.02 | 2780 | 343.35 |
| 7 | 2.584 | 19055706 | 97.22 | 3151008 | 432.06 |
| 8 | 3.132 | 448622 | 2.29 | 59232 | 693.03 |
| 9 | 4.543 | 1040 | 0.01 | 132 | 416.46 |

**Supplementary Figure 6. LC-MS data for repurchased compound 5.**

[6]

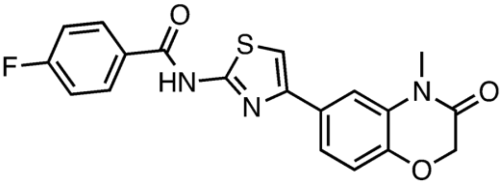


Chemical Formula: $C_{19}H_{14}FN_3O_3S$
Exact Mass: 383.07

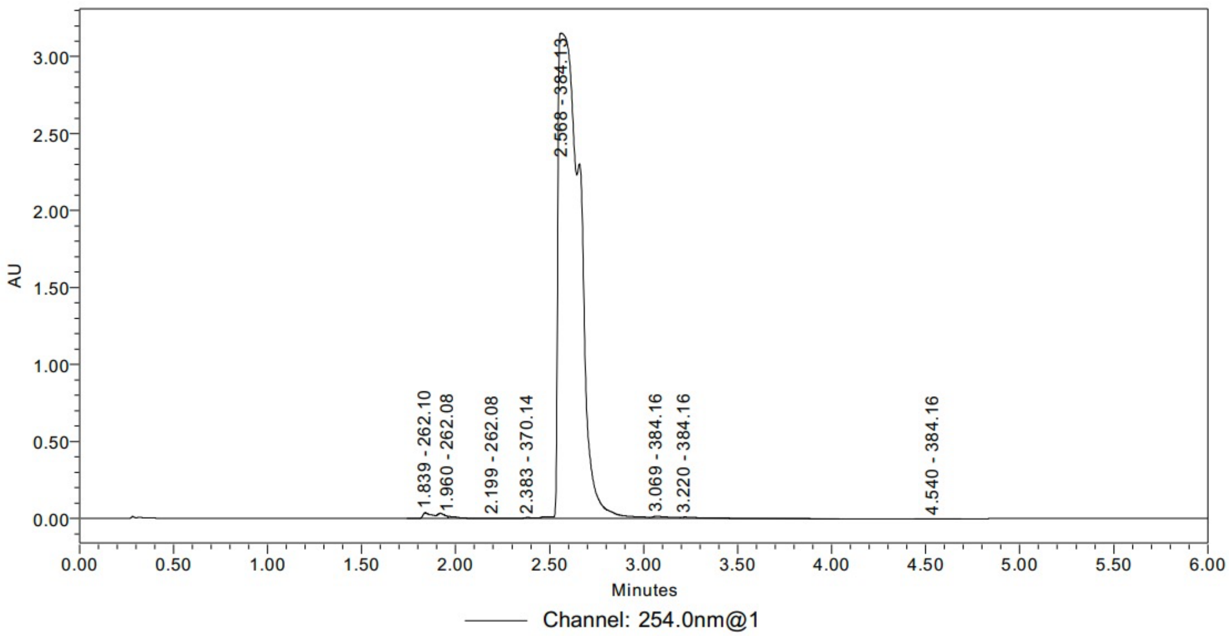


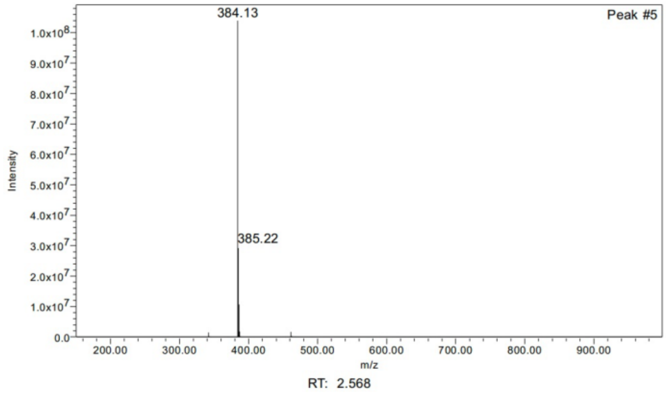


| | RT | Area | % Area | Height | Base Peak (m/z) |
|---|---|---|---|---|---|
| 1 | 1.839 | 213095 | 0.82 | 37046 | 262.10 |
| 2 | 1.960 | 54540 | 0.21 | 14755 | 262.08 |
| 3 | 2.199 | 14344 | 0.06 | 1984 | 262.08 |
| 4 | 2.383 | 31357 | 0.12 | 7191 | 370.14 |
| 5 | 2.568 | 25396304 | 97.86 | 3150257 | 384.13 |
| 6 | 3.069 | 106614 | 0.41 | 15383 | 384.16 |
| 7 | 3.220 | 134018 | 0.52 | 10417 | 384.16 |
| 8 | 4.540 | 774 | 0.00 | 103 | 384.16 |

**Supplementary Figure 7. LC-MS data for repurchased compound 6.**

[7]

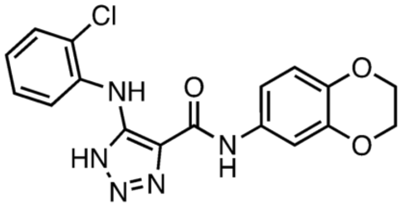


Chemical Formula: $C_{17}H_{14}ClN_5O_3$
Exact Mass: 371.08

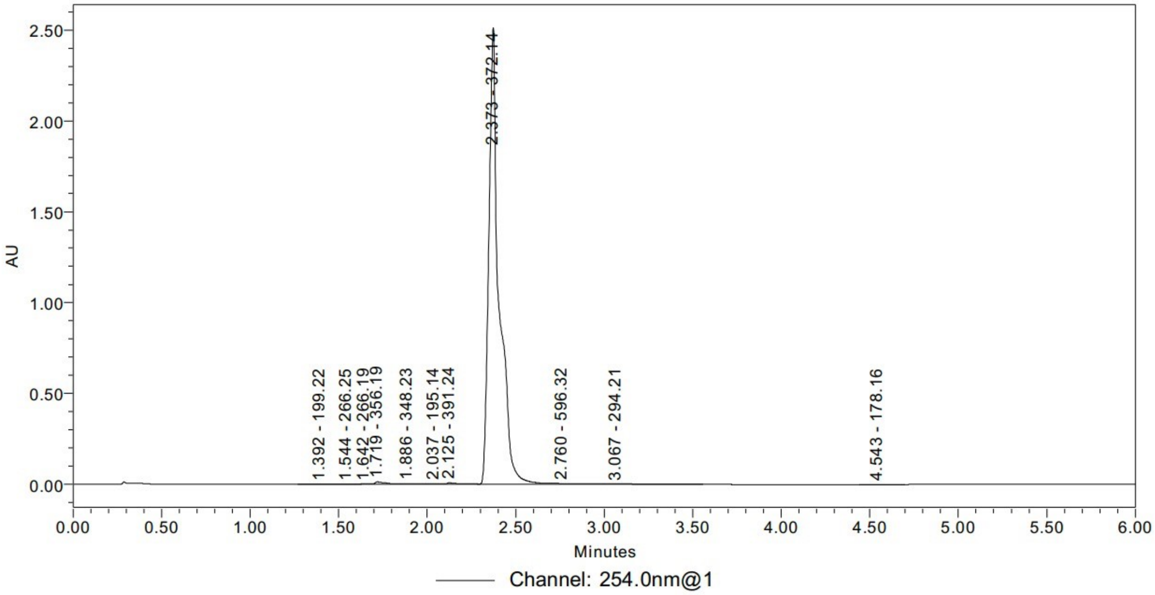


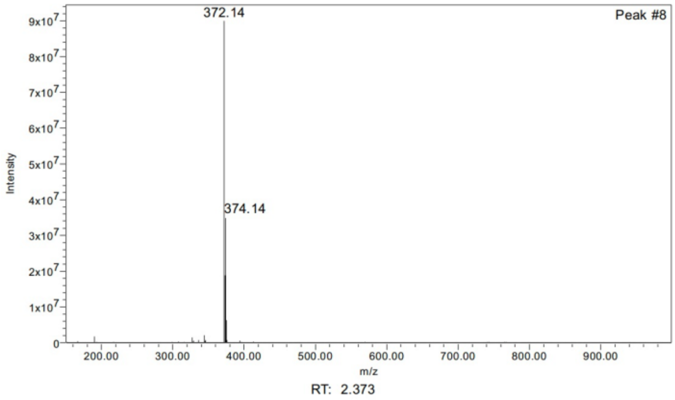


| | RT | Area | % Area | Height | Base Peak (m/z) |
|---|---|---|---|---|---|
| 1 | 1.392 | 4289 | 0.04 | 854 | 199.22 |
| 2 | 1.544 | 4123 | 0.04 | 790 | 266.25 |
| 3 | 1.642 | 6640 | 0.06 | 1736 | 266.19 |
| 4 | 1.719 | 57191 | 0.55 | 13853 | 356.19 |
| 5 | 1.886 | 24455 | 0.23 | 4307 | 348.23 |
| 6 | 2.037 | 19070 | 0.18 | 3894 | 195.14 |
| 7 | 2.125 | 43463 | 0.42 | 9288 | 391.24 |
| 8 | 2.373 | 10199869 | 97.67 | 2513474 | 372.14 |
| 9 | 2.760 | 43973 | 0.42 | 4485 | 596.32 |
| 10 | 3.067 | 38689 | 0.37 | 2848 | 294.21 |
| 11 | 4.543 | 1425 | 0.01 | 171 | 178.16 |

**Supplementary Figure 8. LC-MS data for repurchased compound 7.**

[8]

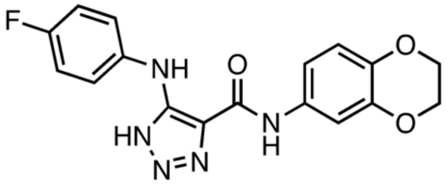

Chemical Formula: $C_{17}H_{14}FN_5O_3$
Exact Mass: 355.11

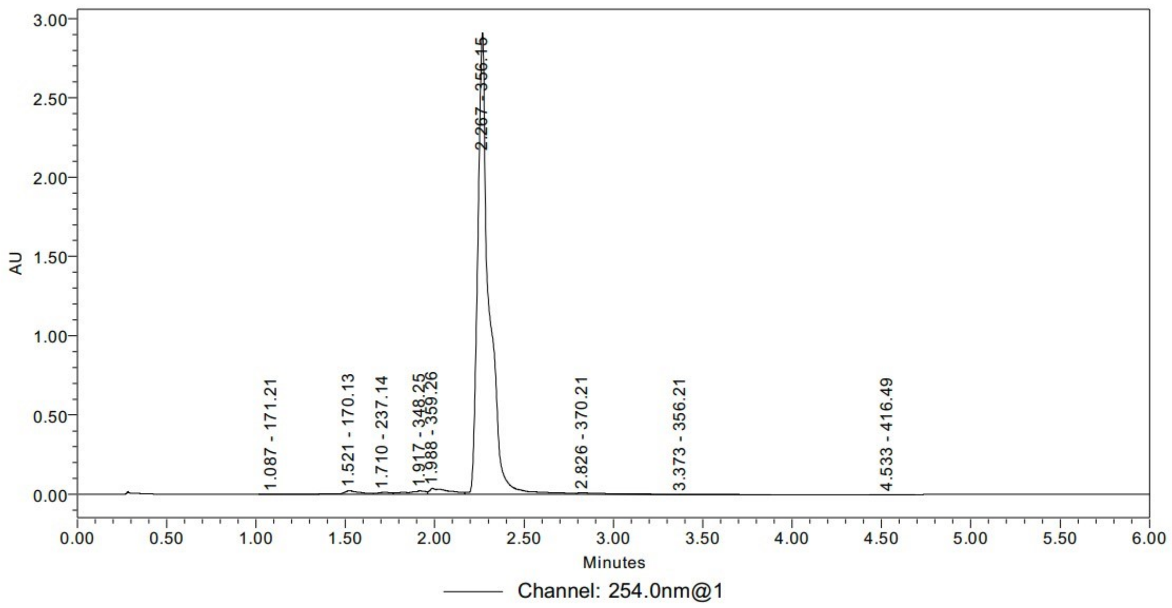


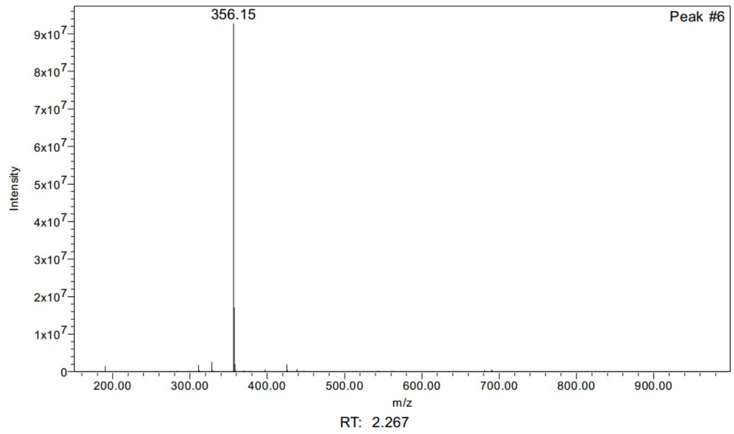


| | RT | Area | % Area | Height | Base Peak (m/z) |
|---|---|---|---|---|---|
| 1 | 1.087 | 1220 | 0.01 | 261 | 171.21 |
| 2 | 1.521 | 161194 | 1.18 | 23665 | 170.13 |
| 3 | 1.710 | 66372 | 0.48 | 11467 | 237.14 |
| 4 | 1.917 | 165063 | 1.20 | 21472 | 348.25 |
| 5 | 1.988 | 287874 | 2.10 | 38028 | 359.26 |
| 6 | 2.267 | 12837925 | 93.61 | 2912701 | 356.15 |
| 7 | 2.826 | 171891 | 1.25 | 10944 | 370.21 |
| 8 | 3.373 | 20968 | 0.15 | 1955 | 356.21 |
| 9 | 4.533 | 2093 | 0.02 | 253 | 416.49 |

**Supplementary Figure 9. LC-MS data for repurchased compound 8.**

[9]

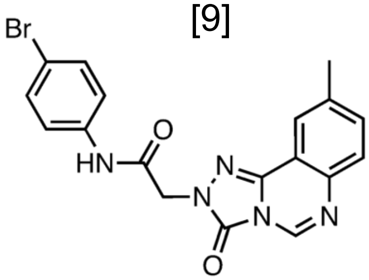


Chemical Formula: $C_{18}H_{14}BrN_5O_2$
Exact Mass: 411.03

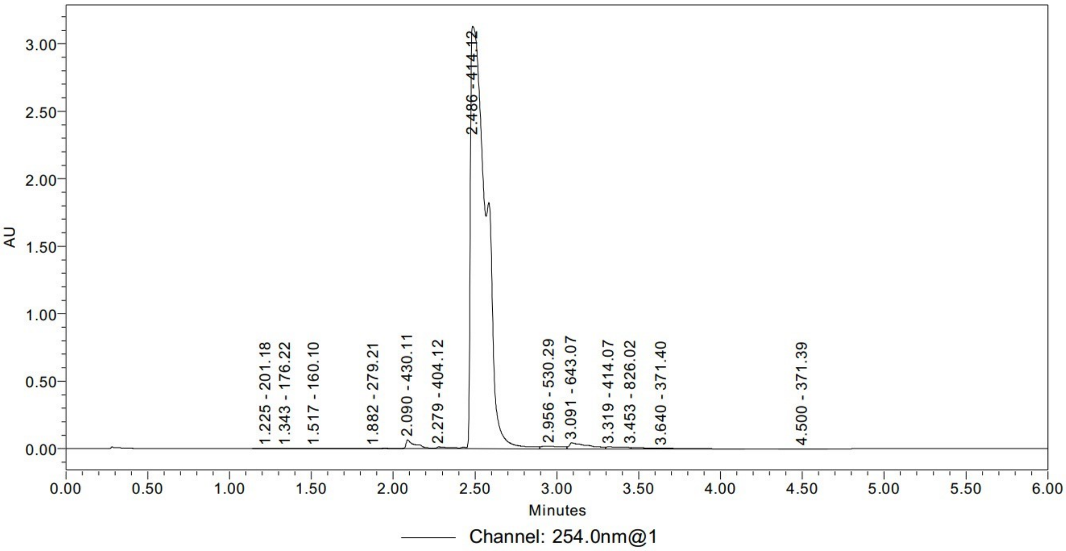


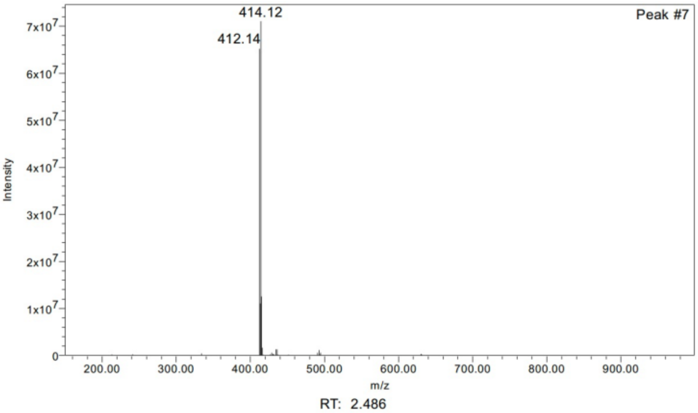


| | RT | Area | % Area | Height | Base Peak (m/z) |
|---|---|---|---|---|---|
| 1 | 1.225 | 4785 | 0.02 | 1135 | 201.18 |
| 2 | 1.343 | 2520 | 0.01 | 588 | 176.22 |
| 3 | 1.517 | 18676 | 0.08 | 1928 | 160.10 |
| 4 | 1.882 | 24265 | 0.11 | 2356 | 279.21 |
| 5 | 2.090 | 289650 | 1.31 | 65699 | 430.11 |
| 6 | 2.279 | 80082 | 0.36 | 13442 | 404.12 |
| 7 | 2.486 | 20910670 | 94.62 | 3130827 | 414.12 |
| 8 | 2.956 | 168479 | 0.76 | 19501 | 530.29 |
| 9 | 3.091 | 349696 | 1.58 | 44182 | 643.07 |
| 10 | 3.319 | 114914 | 0.52 | 13887 | 414.07 |
| 11 | 3.453 | 81792 | 0.37 | 9853 | 826.02 |

**Supplementary Figure 10. LC-MS data for repurchased compound 9.**

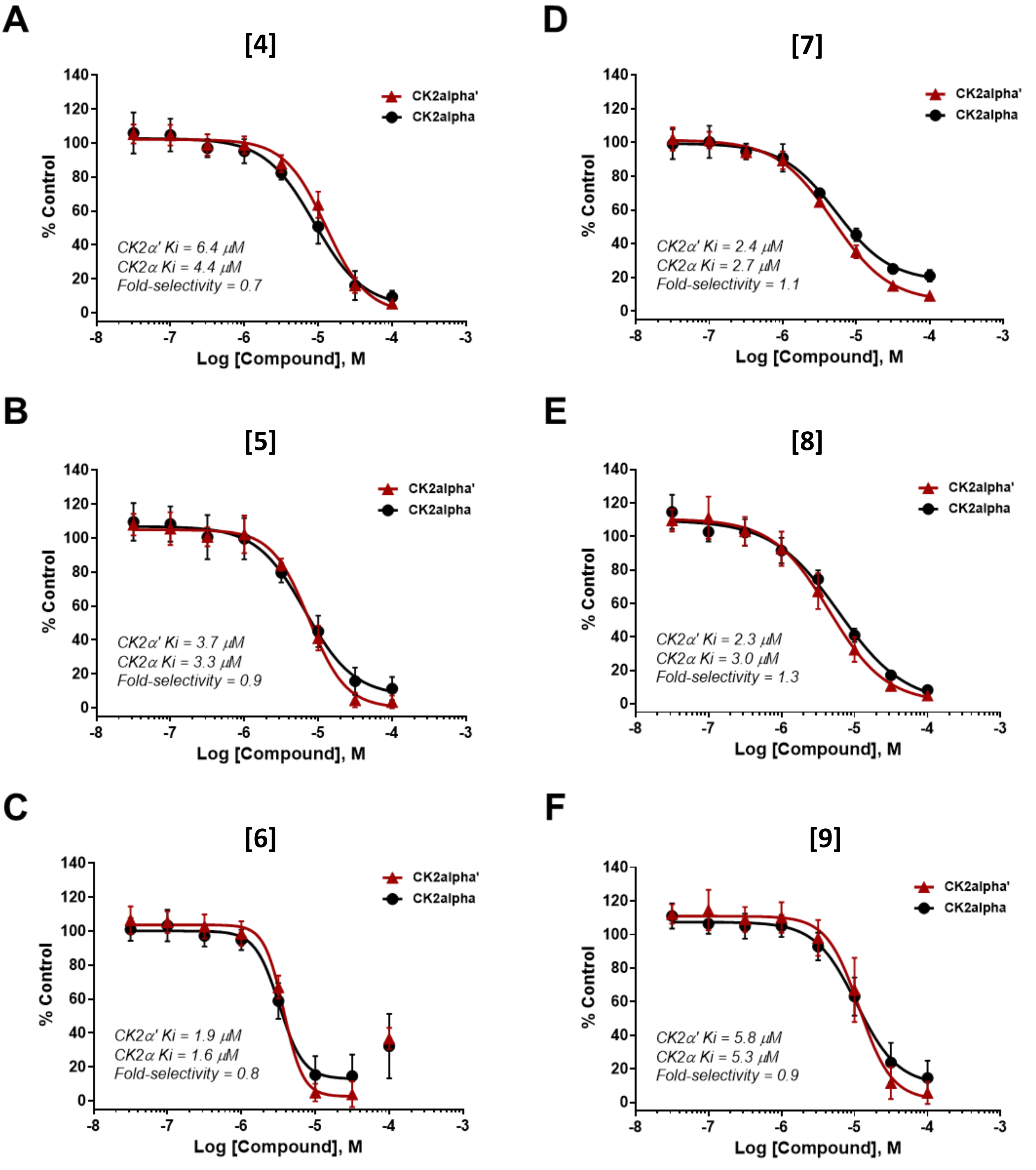


**Supplementary Figure 11.** Dose response curves of six (compounds 4-9) of the seven repurchased hits (A-F) that showed no selectivity between CK2α' and CK2α.

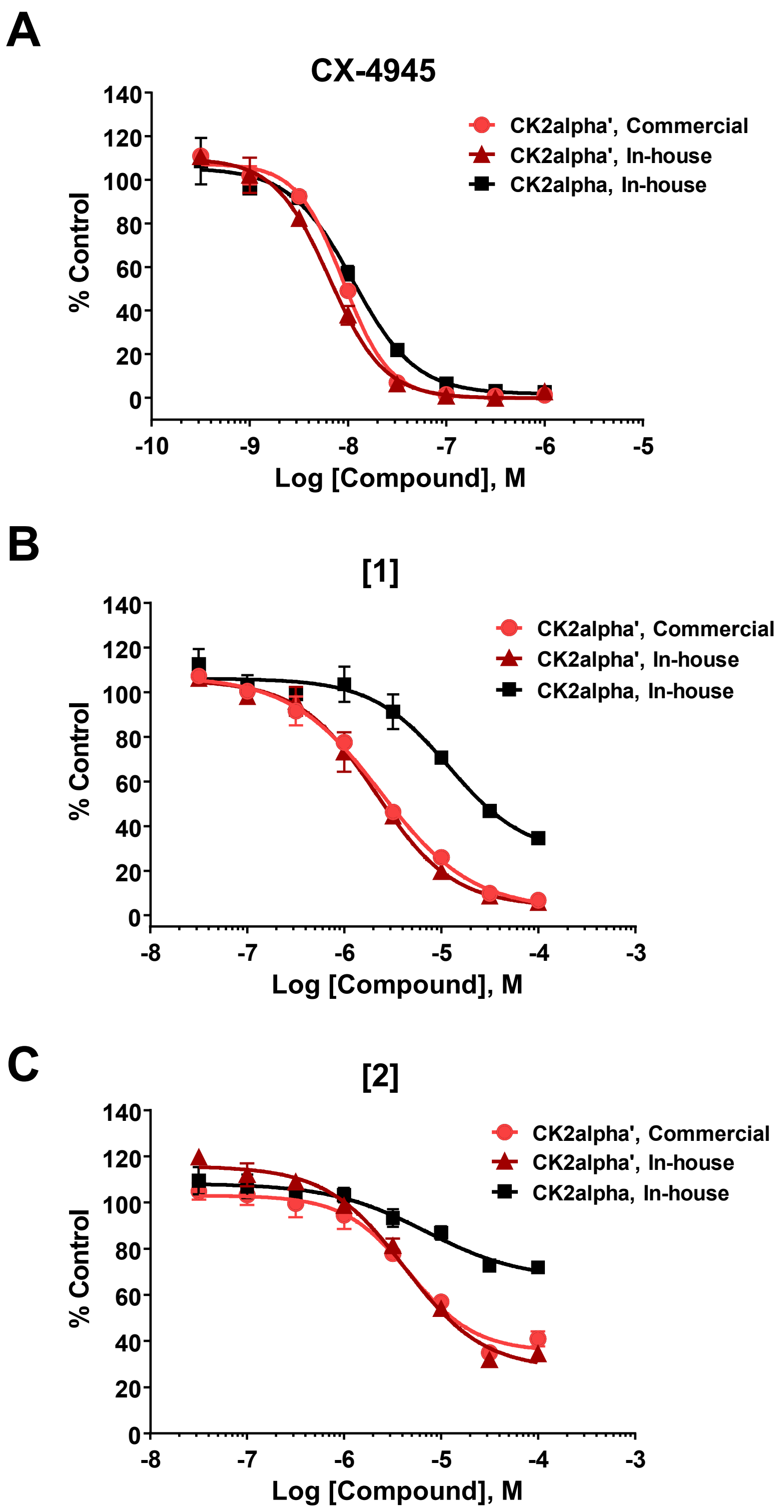


**Supplementary Fig. 12.** Dose response curves of CX-4945, 1 and 2 using the CK2α'-GST enzyme (produced in-house and used in the HTS) and a commercially obtained His-tagged CK2α' enzyme are nearly identical, demonstrating that the GST tag used in the in-house produced CK2α' is not responsible for introducing the observed selectivity over CK2α.

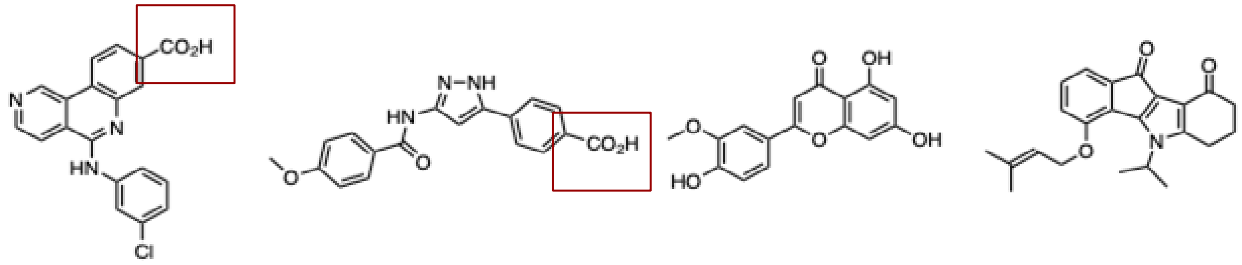

**CX-4945** **15** **16** **17**

**Supplementary Fig. 13. Comparison between CX-4945 and previously reported CK2α' selective inhibitors**. CX-4945 ($IC_{50}$(μM) 0.001), non-selective, **15**; CK2α' $IC_{50}$(μM) 0.14, 2-fold selective [Hou et al.]), (**16;** CK2α' $IC_{50}$(μM) 0.06, 7-fold selective [Baier et al.]), and (**17**; CK2α' $IC_{50}$(μM) 0.273, 2-fold selective [ Lindenblatt et al.]). Red boxes highlight the carboxylic acid moiety.
